# ULK1 forms distinct oligomeric states and nanoscopic structures during autophagy initiation

**DOI:** 10.1101/2020.07.03.187336

**Authors:** Chiranjib Banerjee, Dushyant Mehra, Daihyun Song, Angel Mancebo, Do-Hyung Kim, Elias M. Puchner

**Affiliations:** School of Physics and Astronomy, University of Minnesota, Twin Cities; Department of Biochemistry, Molecular Biology, and Biophysics, University of Minnesota, Twin Cities; Department of Biomedical Engineering and Physiology, Mayo Clinic, Rochester, MN

**Keywords:** Autophagy, ULK1, endogenous tagging, PALM, single molecule localization microscopy, super-resolution, co-localization

## Abstract

Autophagy induction involves extensive molecular and membrane reorganization. Despite significant progress, the mechanism underlying autophagy initiation remains poorly understood. Here, we used quantitative Photoactivated Localization Microscopy with single-molecule sensitivity to analyze the nanoscopic distribution of endogenous ULK1, the kinase that triggers autophagy. Under amino acid starvation, ULK1 formed large clusters containing up to 161 molecules at the endoplasmic reticulum. Cross-correlation analysis revealed that ULK1 clusters engaging in autophagosome formation require 30 or more molecules. The ULK1 structures with more than the threshold number contained varying levels of Atg13, Atg14, Atg16, LC3B, GEC1 and WIPI2. We found that ULK1 activity is dispensable for the initial clustering of ULK1, but necessary for the subsequent expansion of the clusters, which involves interaction with Atg14, Atg16, and LC3B and relies on Vps34 activity. This quantitative analysis at the single molecule level has provided unprecedented insights into the behavior of ULK1 during autophagy initiation.

## Introduction

Macroautophagy (hereafter referred to as autophagy) is an evolutionarily conserved process in eukaryotic cells that degrades intracellular organelles and macromolecules in the lysosome. Autophagy plays a fundamental role in maintaining the cellular homeostasis via controlling degradation and recycling of cellular constituents. Dys-regulation of autophagy is related to a broad range of diseases, including cancer, neurodegeneration, and diabetes^1, 2^. Autophagy is mainly induced under amino acid starvation or other stress conditions via suppression of mechanistic target of rapamycin complex 1 (mTORC1), the central nutrient-sensitive protein kinase complex and subsequent activation of UNC51-like kinase 1 (ULK1), the protein kinase that triggers molecule events necessary for autophagy initiation ^3, 4, 5, 6, 7, 8^. Despite the extensive studies on autophagy mechanisms, our understanding of the autophagy initiation process remains still limited largely due to the complex nature of the process.

Autophagy is an extensive membrane process that starts with the formation of the double membrane structure called phagophore. The phagophore membrane grows and develops into the closed double membrane structure called autophagosome that encompasses cargos destined to the lysosome^9, 10, 11^. According to the prevalent model, autophagosome formation starts at the endoplasmic reticulum (ER) exit sites or the ER-mitochondria contact sites with the formation of the phagophore. The phagophore membrane has also been shown to originate from late endosomes or lysosomes, the Golgi complex as well as mitochondria^9, 10, 11^. The key step for autophagosome formation is activation of ULK1^5, 6, 7, 8^. ULK1 forms a large protein complex by interacting with Atg13, FIP200 (FAK family kinase interacting protein 200 kDa), and Atg101^5, 6, 7, 8, 12, 13, 14, 15^. ULK1 mostly localizes on the ER or Atg9 compartments^16^ and forms puncta in the cytoplasm of starved cells prior to the formation of the phagophore^17, 18, 19^. This suggests that understanding how ULK1 puncta form in response to starvation is key to better understand the autophagy initiation process.

Prior studies employing confocal or other conventional fluorescence microscopy techniques have provided useful but limited knowledge on the nature of ULK1 puncta due to their limited spatial resolution and lack of single molecule sensitivity. Recently, highly sensitive single-molecule and super-resolution microscopy techniques have been developed, enabling the study of autophagy below the optical diffraction limit^20^. For instance, Structured Illumination Microscopy (SIM) is a fast technique for imaging living cells, achieves a lateral resolution of ∼100 nm and yielded significant new insights into autophagy and its initiation process^18, 21, 22, 23^. Recently, Chen et al.^24^ employed SIM to quantify the interaction of lysosomes with mitochondria during mitophagy with increased resolution. Stimulated Emission Depletion (STED) microscopy^25^ offers a higher resolution of ∼50 nm, is also compatible with live cell imaging, but comes at the cost of scanning a high-power excitation and depletion laser beam across the sample. This technique was recently used to study alternative autophagy at the Golgi and enabled to detect translocation of WIPI2 puncta from the cytoplasm to the trans-Golgi^26^. STED also yielded insights into the initiation of autophagosome formation by monitoring LC3B puncta at contact sites between the plasma membrane and the endoplasmic reticulum (ER) contact sites^27^.

The highest resolution of ∼20 nm is offered by Single Molecule Localization Microscopy techniques, but due to the long data acquisition time, they are mostly applied in fixed cells for high-resolution studies. Stochastic Optical Reconstruction Microscopy (STORM)^16, 28^, for instance, has increased our understanding of autophagy and its initiation process due to its capability to differentiate structures with small sizes that cannot be resolved with other techniques. Using STORM, Karanasios et al.^16^ recently resolved small Atg13 clusters and monitored their maturation during autophagy, which clarified the transient participation of Atg9 during autophagy initiation. Photo-activated Localization Microscopy (PALM) has a comparable resolution to STED and STORM but can provide additional information about the stoichiometry of protein complexes or the number of biomolecules in intracellular structures^29, 30, 31, 32, 33, 34, 35^. In particular, quantitative Photoactivated Localization Microscopy (qPALM)^36, 37^ using irreversibly bleaching photoconvertible or photoswitchable fluorescent proteins (PSFPs) can be combined with endogenous tagging e.g. using CRISPR/Cas9 to quantify native molecule numbers in protein clusters or intracellular structures. Various analysis techniques have also been developed for PALM to determine the precise spatial distribution of signaling proteins^38, 39^.

Most fluorescence and super-resolution microscopy studies conducted with mammalian cells have relied on overexpression of fluorescently tagged proteins^40^. It is noteworthy that overexpression can cause artifacts in determining the oligomeric state and spatial distribution of proteins^41^. Many proteins involved in autophagy function as protein complexes. Thus, overexpressing autophagy proteins can disturb the stoichiometry of the complexes as well as the integrity of the autophagy pathway. In addition, overexpression itself can alter the biological function of proteins^42^. A few recent studies have applied PALM with endogenously tagged proteins^43, 44, 45, 46^. However, none of the studies could successfully quantify the number of proteins due to the challenges associated with the blinking behavior of PSFPs.

In this study, we have applied qPALM to analyze the oligomeric states and clustering behavior of endogenous ULK1 with single-molecule sensitivity. Endogenous ULK1 was expressed as a fusion with mEos2, one of the well-characterized PSFPs^32, 47^, via CRISPR-cas9 based genome editing^48^. Under amino acid starvation, ULK1 assembled near the ER forming clusters comprising over 30 ULK1 molecules. We found that the identified threshold number is necessary for ULK1 clusters to participate in autophagosome formation. Through cross-correlation analysis, we determined that all structures containing over 30 ULK1 molecules induced by starvation are associated with Atg13 and Atg14. These structures grew in the size to associate with Atg16, LC3B, GEC1 and WIPI2 for autophagosome maturation. We also determined distinct functions of ULK1 and Vps34 kinase activities in the initial clustering of ULK1 and the subsequent expansion of the clusters.

## Results

### Quantitative analysis of endogenous ULK1 clusters reveals its high-order multimeric states

ULK1 forms puncta in the cytoplasm prior to autophagosome formation^16, 17, 19^. To characterize the properties of ULK1 puncta, we used qPALM and quantified the multimeric states of ULK1 molecules. To avoid potential artifacts of qPALM analysis caused by overexpression (Supplementary Fig. 1), we engineered the HeLa cell genome using the CRISPR/Cas9-assisted genome editing technique^48^ introducing mEos2 into the allele of the ULK1 gene (Fig. 1a). We then selected genome-edited cells that expressed all endogenous ULK1 molecules with mEos2 tagged at the N-terminus (Fig. 1b). The endogenously tagged ULK1 was confirmed to form functional complexes with Atg13 and FIP200 as similarly as untagged endogenous ULK1 (Fig. 1b), and it responded to amino acid starvation for its activation toward phosphorylation of Atg14 Ser29, a ULK1 target site^19^ (Fig. 1c). The genome-edited cells also showed induction of Atg13 and LC3B puncta to a similar extent as unmodified HeLa cells under amino acid starvation (Supplementary Fig. 2). We further confirmed that the endogenous tagging does not affect autophagy flux of HeLa cells as reflected in the differences of LC3B II and p62 levels in the presence and absence of BAFA1, a lysosomal inhibitor (Fig. 1d).

**Fig. 1.**
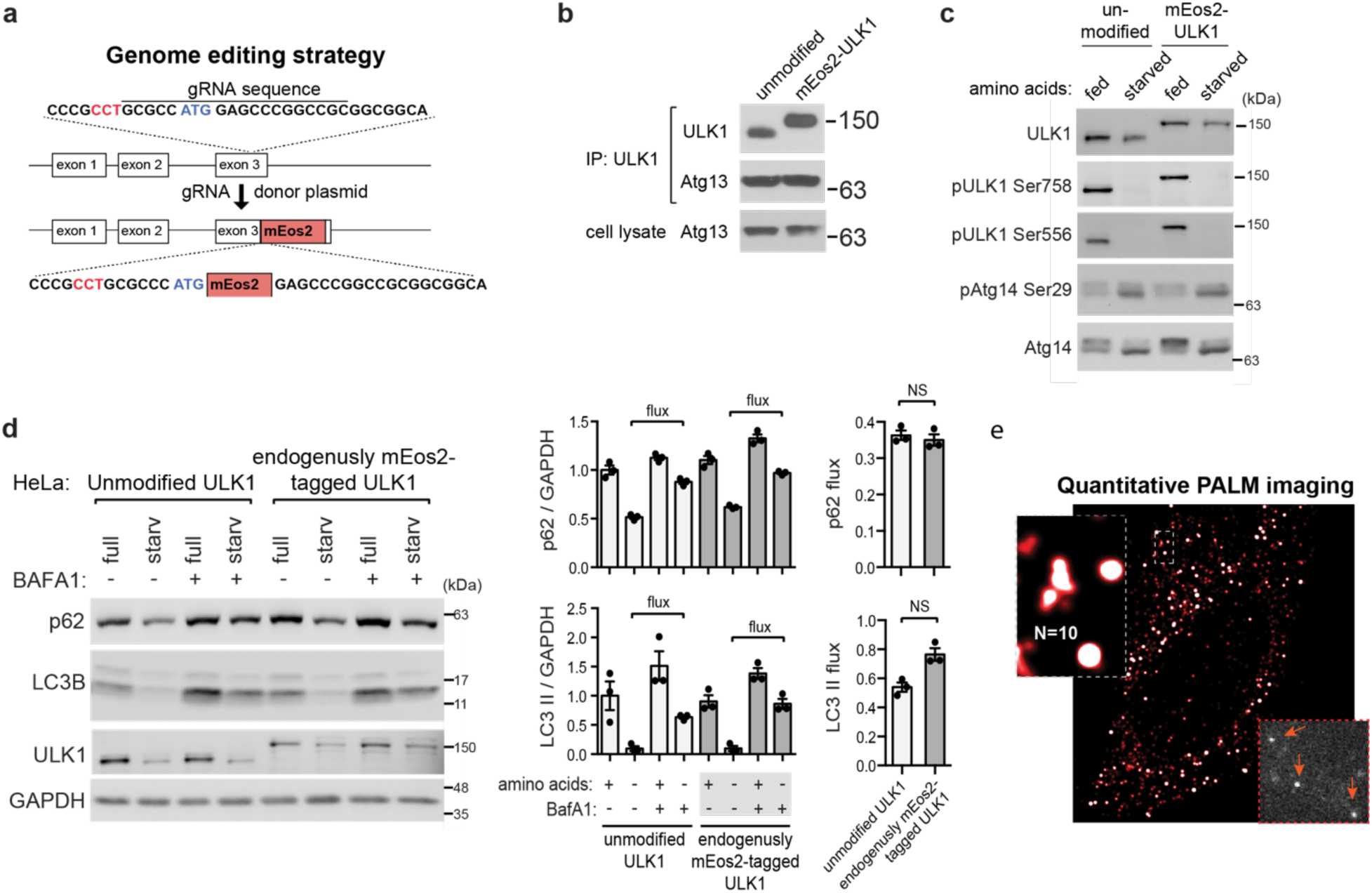
Quantitative PALM reveals the nanoscopic distribution and oligomeric state of endogenous ULK1. **a** Schematics of endogenous tagging of ULK1 with mEos2. The gRNA PAM sequence is depicted in red. The detailed procedure is described in the Method section. **b** The endogenously tagged ULK1 interacts with Atg13 to the same degree as the untagged endogenous ULK1. Immunoprecipitates were obtained from genome-modified HeLa cells (mEos2-ULK1) or unmodified HeLa cells grown in full medium using an anti-ULK1 antibody and analyzed by western blotting for endogenous ULK1 and Atg13. **c** Endogenously tagged ULK1 is functionally intact for activation of ULK1 in response to amino acid starvation. Cells were incubated in amino acid enriched full medium (fed) or amino acid deprived medium (starved) for 60 min. The degree of phosphorylation of Atg14 Ser29, a ULK1 target site^19^, was analyzed by western blotting. **d** Autophagy flux, monitored by the levels of p62 and LC3B II with use of BAFA1, was similar in cells expressing unmodified ULK1 and in those expressing endogenously tagged mEos2-ULK1, as shown by western blot analysis. **e** PALM image (red/white, scale bar 5 µm) showing the nanoscopic distribution and oligomeric state of ULK1 oligomers throughout the cytoplasm. The bottom right inset shows bright single-molecule fluorescence of mEos2-ULK1 in a single data acquisition frame. The zoom on the left exemplifies quantification of oligomeric state of a ULK1 cluster.

Using the genome-edited cells, we conducted PALM analysis of mEos2-ULK1 to determine the oligomeric states of ULK1. Each time a single mEos2 molecule was photoactivated and fluoresced, its center of mass was determined by a Gaussian fit to its intensity profile. Since mEos2, like other fluorophores, can emit a variable number of fluorescent burst that are separated by a variable short dark-time, a single molecule can appear as a cloud of localizations in raw PALM data^29^. To correct for these clustering artifacts, we employed our previously developed method that groups localizations originating from the same mEos2 molecule to a single, photon weighted averaged position based on a spatio-temporal threshold (see Methods and Supplementary Fig. 3)^32^. By applying this blink-correction, we obtained the detected number of ULK1 molecules in ULK1 clusters with ∼25 nm resolution (Supplementary Fig. 3h) as well as the overall distribution of oligomeric states (Fig. 1e). It is important to note that not all mEos2 molecules are detectable as previous studies in different model systems have quantified^32, 33^ (see Methods). Throughout this manuscript we only report the number of detected molecules for clarity.

### Amino acid starvation induces high-order ULK1 oligomers

Regardless of amino acid presence or absence, mEos2-ULK1 was distributed throughout the whole cytoplasm of cells (Fig. 2a). Distinctly in amino acid starved cells, we observed emerging arc-shaped and spherical structures of various sizes (Fig. 2a, right inserts). These structures could not be resolved or discriminated in confocal fluorescence images. Via qPALM analysis, we grouped molecules appearing within a radial distance of 400 nm, the size that can encompass large structures induced by starvation up to a size of 800 nm (Fig. 2a zooms and Methods). In fed and starved conditions, the probability of finding monomeric ULK1 was 30% with the remaining 70% being in higher order oligomeric states. The probability of finding ULK1 clusters containing up to 10 molecules in either fed or starved conditions was 98% (Fig. 2b, insert) corresponding to a fraction of 93% of all ULK1 molecules in fed and 89% in starved cells. Starvation induced 5% of all ULK1 molecules to transition from lower to higher oligomeric states. In fed cells the highest observed number of ULK1 molecules in a structure was 59, whereas in starved cells this number significantly increased to 144. It is important to note that these highest order oligomeric states occur with low probability and are thus not an accurate metric for comparison. Below we quantify with high statistics that the most significant increase in oligomeric states upon autophagy induction by starvation occurs at a threshold number of 30 ULK1 molecules.

**Fig. 2.**
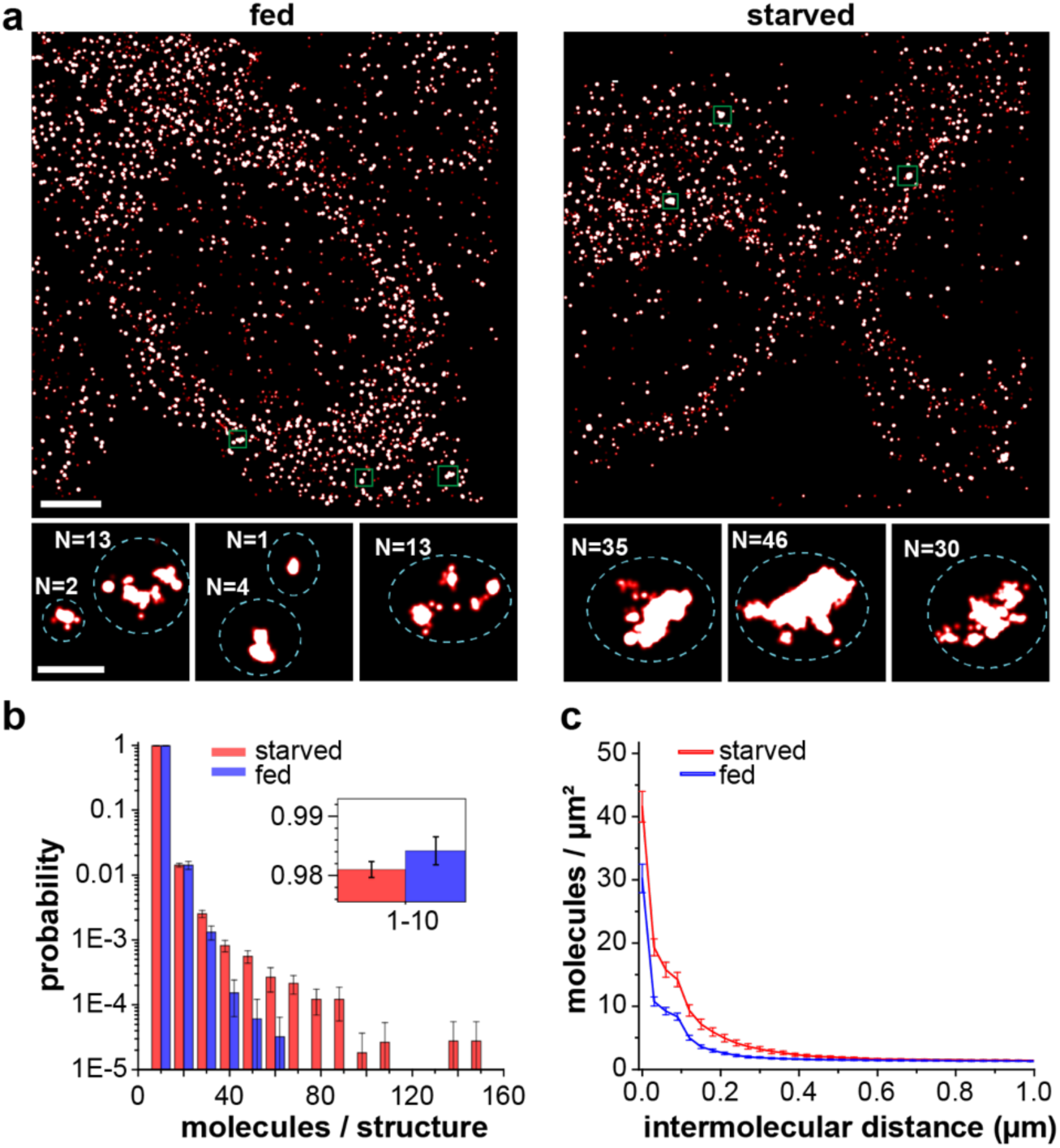
Amino acid starvation induces clusters with more ULK1 molecules and larger size. **a** PALM images of mEos2-ULK1 in amino acid supplemented cells (fed) and amino acid deprived cells (starved) for 150 min. Scale bars: 5 µm. Magnified regions show small diffraction-limited structures, which were impossible to be resolved in conventional fluorescence images, and the corresponding number of ULK1 molecules. Scale bar: 500 nm. **b** The cluster analysis after the blink correction shows that amino acid starvation induced more higher-order oligomers and structures containing up to 144 ULK1 molecules (red) compared to 59 ULK1 molecules in fed cells (blue). The insert shows the top of the first bin to visualize that the majority of ULK1 is in oligomers with 2-10 molecules. Error bars of each bin represent the standard error of the mean (SEM) with a variance calculated from the cell-to-cell variability. **c** The pair correlation function of ULK1 in cells from b quantifies the molecular densities as a function of their intermolecular distances. A peak at short distances reflects the accumulation of molecules with respect to a random distribution. The statistically significant difference between starved (red) and fed cells (blue) at distances between 0 nm and 400 nm reflects an increase of cluster sizes and molecular densities upon starvation. Errors represent the SEM, which is calculated from the values of individual cells (fed: N=21 cells, starved: N=34 cells).

To further clarify how amino acid starvation affects ULK1 clusters, we conducted the pair-correlation analysis. The pair-correlation magnitude, which quantifies the density of molecules as a function of their intermolecular distance, exhibited a peak in starved and fed cells, indicating the accumulation of ULK1 compared to a random distribution. In starved cells, the pair-correlation was significantly higher compared to fed cells at distances from 0 - 600 nm (Fig. 2c). While the absolute difference in the pair correlation magnitude appears to be small, it is important to note that only about 5% of all ULK1 molecules are induced by starvation to form higher-order oligomers. This result corroborates the observation that amino acid starvation induces the formation of larger structures with denser ULK1 molecules^17, 18^.

### Starvation-induced structures with high numbers of ULK1 molecules form within 100 nm of the ER

It is known that autophagy initiation occurs on the surface of the ER^11, 16^. To clarify if our observed starvation-induced structures with large ULK1 molecule numbers form at the ER, we performed co-localization PALM analysis using mEos2-ULK1 HeLa cells transiently transduced with HaloTag-tagged Sec61β, an ER marker protein. The HaloTag protein covalently binds its ligands with an attached organic fluorophore in a one to one ratio producing high photon counts and thus high localization precision^49, 50^. To generate two-color PALM images, we used the JF646 dye, which has no spectral overlap with mEos2. mEos2-ULK1 was localized not only on or near the tubular ER regions but also to some extent throughout the cytoplasm in both fed and starved cells (Fig. 3a). In starved cells, the induced structures with many ULK1 molecules were found in contact or close to the ER (Fig. 3a, lower right and Supplementary Fig. 4). In order to determine the location of starvation-induced structures containing many ULK1 molecules with respect to the ER, we separately analyzed ULK1 molecules that appeared within a 100 nm distance from an ER localization (see Methods). In fed cells, the distribution of the number of ULK1 molecules and the radii of structures were similar whether they were in proximity to the ER or not (Fig. 3b, left). Only in starved cells, we again observed structures with a large number of ULK1 molecules that occurred at the ER and were not present in fed cells.

**Fig. 3.**
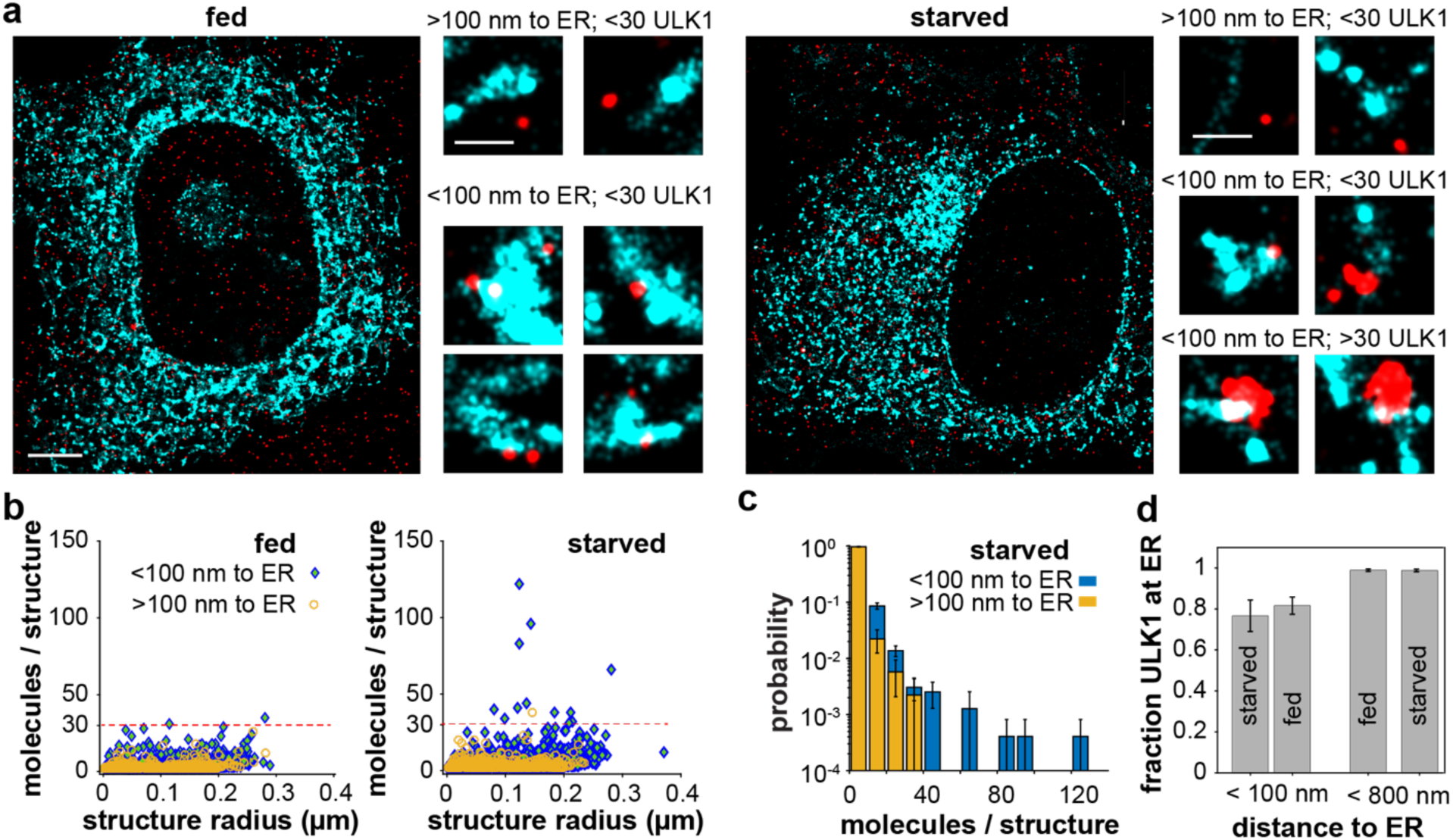
Starvation induced ULK1 structures are in proximity to the ER. **a** Two-color PALM images of mEos2-ULK1 (red) and the ER marker HaloTag-Sec61β (cyan) in fed (left) and starved cells (right). Zoom images of ULK1 and the ER show the presence of ULK1 clusters only on or near the ER in fed cells. In starved cells larger ULK1 structures with more than 30 ULK1 molecules are in contact or in proximity to the ER (lower right). Main scale bar: 5 µm, zoom scale bar: 1 µm. **b** Quantification of the number of ULK1 molecules and the approximated radius of structures closer than 100 nm (blue) and further away from the ER (yellow) in fed and starved cells. Structures with a large number of ULK1 molecules that are later identified to contain Atg13 are only found in close proximity to the ER in starved cells. **c** Normalized histogram of the number of ULK1 molecules in structures of starved cells. Structures in proximity to the ER exhibit the highest number of ULK1 molecules. **d** Fraction of ULK1 molecules at a given distance from the ER in fed and starved cells. In both fed and starved cells, about 80% of all ULK1 molecules were within a distance of 100 nm from an ER localization whereas over 95% of ULK1 molecules were within a distance of 800 nm. Errors represent the SEM from four fed and five starved cells.

To further visualize the distinct population of structures with a high oligomeric state of ULK1 at the ER, we created a normalized histogram of the number of ULK1 molecules in each structure (Fig. 3c). Structures that had a distance of more than 100 nm from an ER localization contained up to 38 ULK1 molecules, whereas structures closer than 100 nm contained up to 122 ULK1 molecules. Therefore, starvation induced structures that contain a high number of ULK1 molecules are located at and near the ER. In both fed and starved cells, about 80% of all ULK1 molecules were within a distance of 100 nm from an ER localization whereas over 95% of ULK1 molecules were within a distance of 800 nm (Fig. 3d). No significant difference of the ULK1-localization with respect to the ER was observed between fed and starved cells. These results suggest that ULK1 molecules engaging in autophagy are localized at and near the ER, where they accumulate to form larger structures with a large number of ULK1 molecules.

### ULK1 structures containing Atg13 exhibit considerable variability in size, shape and multimeric state

ULK1 interacts with Atg13 to regulate autophagy^5, 6, 7^. We therefore wondered whether the higher-order multimeric states of ULK1 in starved cells are related to the autophagy initiation complex containing both ULK1 and Atg13. To address this, we performed co-localization PALM experiments monitoring both ULK1 and Atg13. We transiently expressed Atg13 as a fusion with HaloTag in the mEos2-ULK1 HeLa cells and treated the cells with the HaloTag dye-ligand JF646. In fed cells, both Atg13 and ULK1 formed small clusters that did or did not co-localize with each other throughout the cytoplasm (Fig 4a, left). Starved cells also contained small ULK1 clusters, many of which did not colocalize with Atg13. However, starvation induced large arc-shaped and spherical structures that contained both ULK1 and Atg13 (Fig. 4a, right and Supplementary Fig. 5). These structures were not observed in fed cells. ULK1 structures associated with Atg13 exhibited a variety of sizes, shapes, and multimeric states. Some of the structures contained small, dense ULK1 clusters, whereas some contained large ULK1 clusters with various oligomeric states (Fig. 4b). In contrast, ULK1 molecules that did not colocalize with Atg13 only formed small clusters (Fig. 4b, right). This co-localization PALM analysis showed that the clustering pattern is different between ULK1 structures that colocalized with Atg13 and those that did not colocalize with Atg13.

**Fig. 4.**
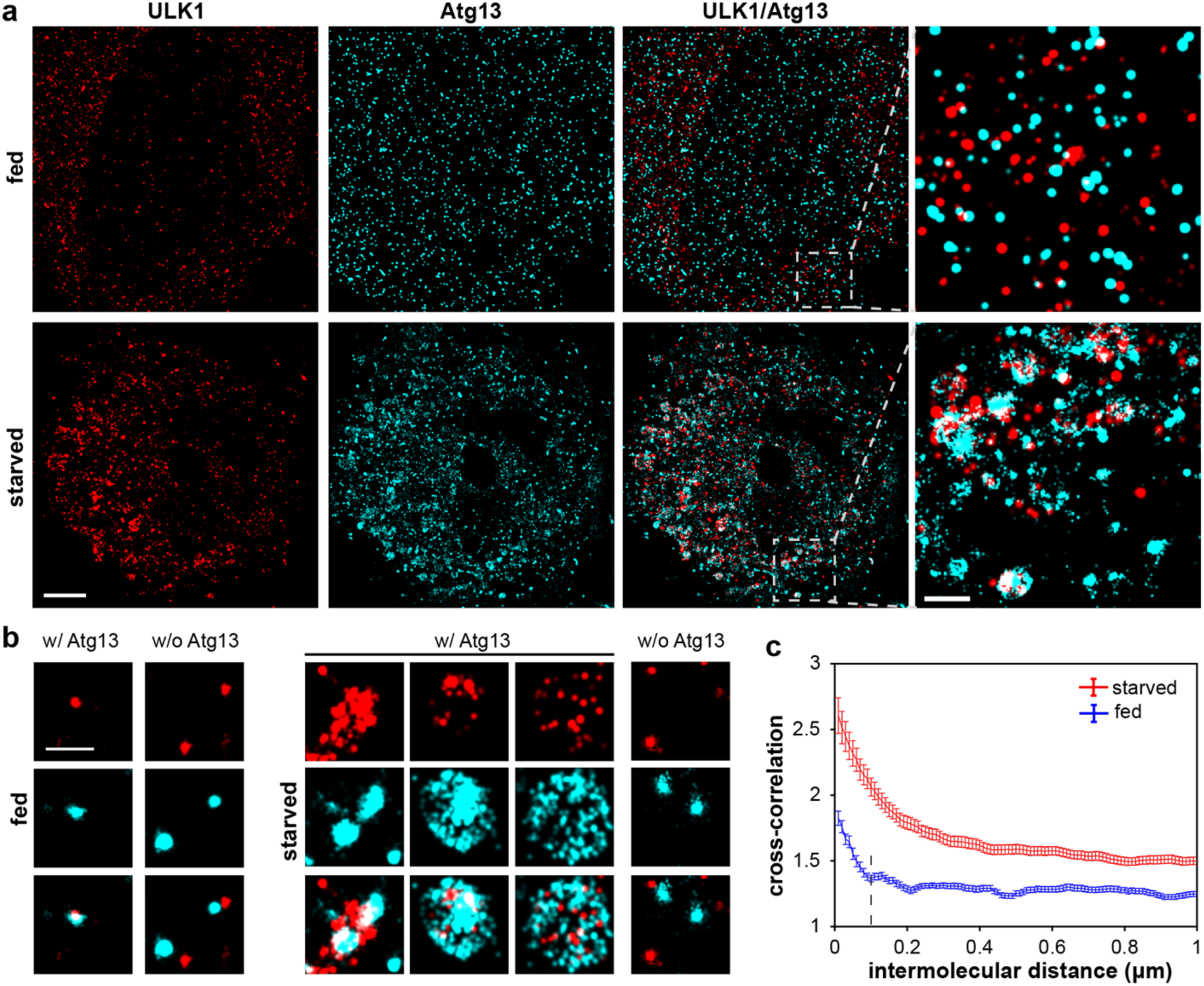
ULK1 clusters containing Atg13 display variability in size, shape, and multimeric state under starvation. **a** Left: Two-color overlay of PALM images of mEos2-ULK1 (red) and HaloTag-Atg13 bound to JF646 (cyan) in fed (top) and starved cells (bottom). Scale bar: 5 µm. Right: Magnified images show the presence of various small dense ULK1 and Atg13 clusters that do or do not co-localize (upper). Upon starvation, larger spherical and arc-shaped structures containing both Atg13 and ULK1 become visible (lower). Scale bar: 1 µm. **b** Left: Examples of magnified two-color PALM images in fed cells show small ULK1 cluster that do not co-localize with Atg13. Right: In starved cells Atg13 forms denser and larger arc-shaped or spherical structures that are associated with ULK1. ULK1 that does not contain Atg13 only forms small cluster. Scale bar: 500 nm. **c** Cross-correlation of ULK1 and Atg13 shows colocalization up to a distance of 100 nm between ULK1 and Atg13 in fed cells (blue) and a significantly larger degree of co-localization beyond 200 nm in starved cells (red). The error bar corresponds to SEM from seven fed and ten starved cells. Condition: 150 min of amino acid starvation with 180 nM bafilomycin A1 (BAFA1) to increase the number of detected autophagosomes in each cell by inhibiting the fusion of autophagosomes with lysosomes.

To further clarify the nature of ULK1 clusters that colocalize with Atg13, we performed cross-correlation analysis^39, 51^ of ULK1 and Atg13. Like pair correlation analysis, cross-correlation analysis of two-color PALM data allows for robust quantification of the co-distribution of two different proteins^45, 51^. When the two proteins are randomly distributed with respect to each other, the cross-correlation value becomes one. However, when two proteins interact and are more likely to co-localize within a short distance, the magnitude of the cross-correlation is larger than one up to that distance. Using the cross-correlation analysis, we measured the extent of co-localization between ATG13 and ULK1. In fed cells, we observed a peak of the cross-correlation at short distances up to 100 nm; however, under starvation the peak was significantly higher in a broad range of distance (Fig. 4c). This suggests a substantially increased level of co-localization between ULK1 and Atg13 within enlarged structures induced by starvation, which might be relevant to autophagosome formation. In our previous report^6^, we used co-immunoprecipitation analysis to investigate the ULK1-Atg13 interaction in response to amino acid starvation, but no significant difference was detected. However, this method has limitations in capturing the dynamics of protein interactions in cells, as protein-protein interactions are often disrupted in vitro during the co-immunoprecipitation process. In contrast, our current study demonstrates, through co-localization analysis, that ULK1 and Atg13 interact dynamically and form multimeric structures during the initiation of autophagy under amino acid starvation.

### Starvation induces the formation of distinct Atg13-containing structures with over 30 ULK1 molecules

To clarify which of the observed structures are involved in autophagy, we separately analyzed the ULK1 molecules that have a distance shorter than 100 nm to Atg13 (Atg13-bound structures) and those that have a distance longer than 100 nm (Atg13-free structures) (Fig. 5a and Methods). As in the previous quantification of the oligomeric state of ULK1 in proximity to the ER (Fig. 3), we determined the number of ULK1 molecules in each structure and in addition approximated their radius (Fig. 5b and Methods). In fed cells, ULK1 showed similar distribution patterns regardless of Atg13-bound states with a maximum number of 43 ULK1 molecules (Fig. 5b). There was also no detectable difference in the radii of the two types of ULK1 structures. In contrast, amino acid starvation induced significant differences between the two types of ULK1 structures. Atg13-free structures had no detectable difference in their number of ULK1 molecules and in their radii compared to fed cells, whereas Atg13-bound structures contained significantly more ULK1 molecules up to 161 under starvation (Fig. 5b, right). By comparing the abundance of structures that have a large number of ULK1 molecules in fed and starved cells, we identified a threshold number of 30 ULK1 molecules. At this threshold, the difference in the distribution of the number of structures was most significant between fed and starved cells by various metrics (Supplementary Fig. 6). In short, if a lower threshold is set, the difference of Atg13-bound structures between starved and fed cells becomes smaller. If a larger threshold is set, the difference of Atg13-bound structures between starved and fed cells also becomes smaller and in addition, the number of structures in fed cells becomes too small and increases the statistical uncertainty of the comparison.

**Fig. 5.**
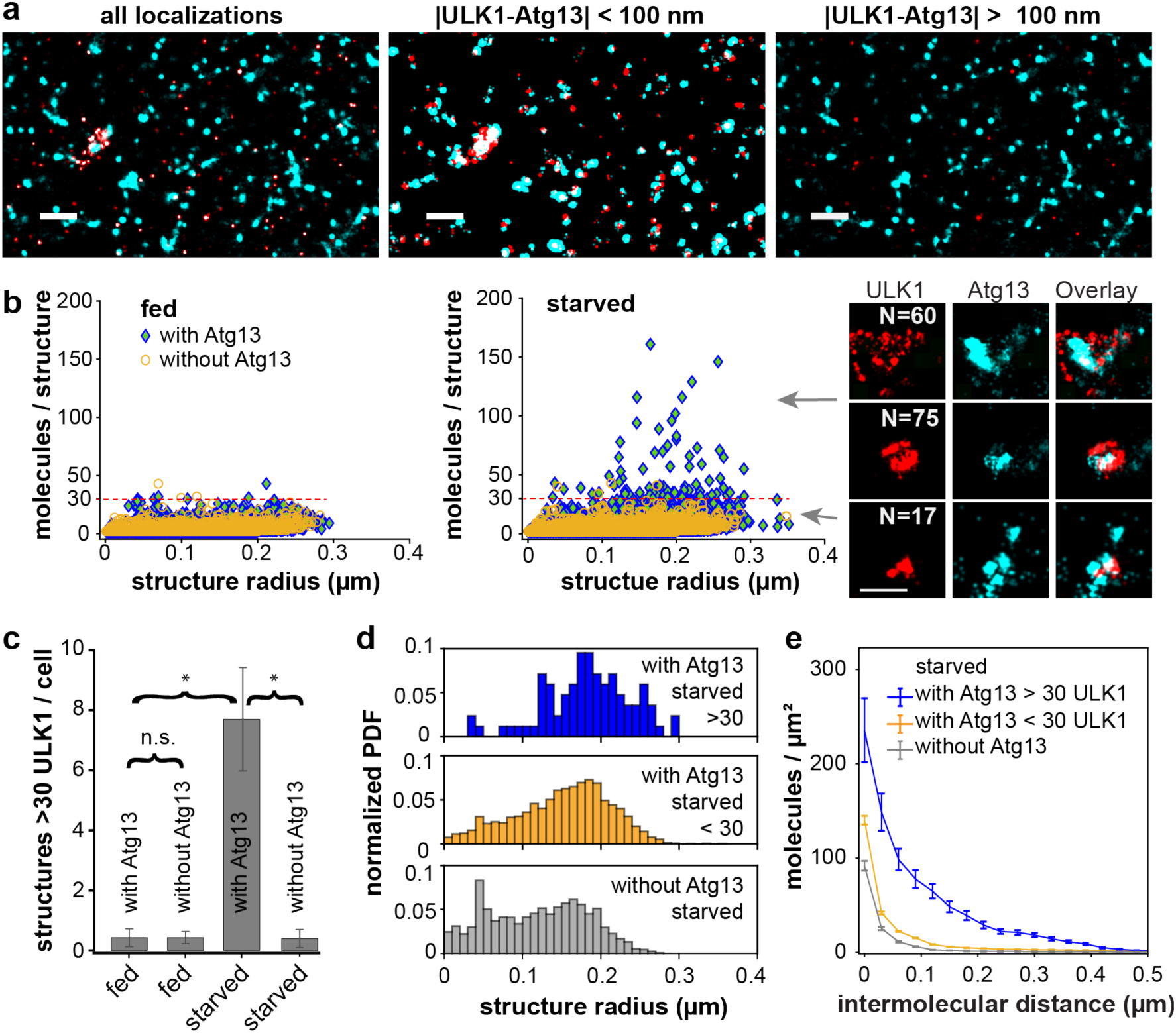
Cross-correlation analysis of two-color PALM data identifies starvation-induced ULK1 structures containing Atg13 and more than 30 ULK1 molecules. **a** Left: Two-color PALM image of all mEos2-ULK1 (red) and Atg13-HaloTag JF646 (cyan) in starved cells. Middle: ULK1 molecules and clusters within 100 nm separation of Atg13 form highly dense and larger arc- or spherical structures. Right: ULK1 separated by more than 100 nm from Atg13 only forms small clusters. Scale bar: 1 µm. **b** Quantification of the number of molecules and radius of ULK1 associated structures. ULK1 structures within a 100 nm distance of Atg13 (with Atg13, blue) were compared to ULK1 structures not in proximity to Atg13 (without Atg13, orange). Left: ULK1 structures exhibited similar distributions in fed cells regardless of Atg13-bound states. Middle: Small, dense clusters as well as larger structures with more than 30 up to 161 ULK1 molecules form under starvation only when they contain Atg13. The dashed line at 30 molecules indicates a threshold used for further analysis. Right: the inset images exemplify large Atg13-containing structures as well as a small dense cluster with less ULK1 molecules under starvation. Scale bar: 500 nm. **c** Number of structures with more than 30 ULK1 molecules per cell in different conditions. In fed cells, there was no significant difference between the number of structures with or without Atg13 per cell, whereas a significant difference was shown with starved cells (ANOVA, p=0.05). Error bars are SEM. **d** Probability density function (PDF) of the radii of structures with more than 30 ULK1 molecules with Atg13, with more than 30 ULK1 molecules without Atg13 (middle), and all other ULK1 molecules without Atg13. **e** Pair-correlation quantifies the density of ULK1 molecules as a function of their distance on the three different types of ULK1 structures. Data was recorded from seven fed and ten starved cells. Condition: 150 min of amino acid starvation with 180 nM bafilomycin A1 (BAFA1) to increase the number of detected autophagosomes in each cell by inhibiting the fusion of autophagosomes with lysosomes.

Starved cells had on average about 7-8 structures per cell that contained both Atg13 and more than 30 ULK1 molecules, which is an about 18-fold increase compared to fed cells (Fig 5c). It is noteworthy that we used the lysosomal inhibitor BAFA1 only in these experiments to increase the statistics for an accurate determination of the threshold of 30 ULK1 molecules. The number of structures with more than 30 ULK1 molecules also significantly increased over time after starvation even in the absence of BAFA1 (Supplemental Fig. 7).

The normalized histogram of the radii of Atg13-free structures was roughly constant up to a radius of ∼200 nm, whereas Atg13-bound structures showed different distributions depending on the number of ULK1 molecules (Fig. 5d). When Atg13-bound structures contained less than 30 ULK1 molecules, the radii spanned a broad range from the smallest measurable values up to 300 nm and exhibited a peak at approximately 200 nm (Fig. 5d). When Atg13-bound structures contained more than 30 ULK1 molecules, most radii were larger than 100 nm and radii up to 300 nm were detected. These results suggest that Atg13-bound structures with more than 30 ULK1 molecules tend to be larger compared to structures with less than 30 ULK1 molecules or structures without Atg13. To further characterize the density of ULK1 in these three classes of structures, we calculated the pair-correlation function, which quantifies the average density of neighboring ULK1 molecules relative to each ULK1 molecule at a given distance. Atg13-free structures formed smallest clusters with a low ULK1 density, whereas Atg 13 bound structure with less than 30 molecules had slightly higher density and size. In contrast, Atg13-bound structures with more than 30 ULK1 molecules formed the densest and by far the largest structures (Fig. 5e). All but one of these structures with more than 30 ULK1 molecules were located within a 100 nm distance from the ER as shown above (Fig. 3b, right and Supplementary Fig. 4).

In summary, our analysis identified unique starvation induced structures that contained more than 30 ULK1 molecules as well as Atg13. These structures were not found in fed cells. While only about 4% of all ULK1 molecules were found in these structures in starved cells, the abundance of these structures in starved cells was significantly higher (∼18 fold) compared to fed cells. ULK1 clusters that did not associate with Atg13 had a lower density and were smaller compared to Atg13-bound structures with more than 30 ULK1 molecules, which highlights the different state of ULK1 when associated with Atg13. We speculate that those Atg13-free ULK1 molecules might function in non-autophagic processes, such as ER-to-Golgi trafficking^52^.

### A quantitative co-localization map of ULK1 with other autophagy proteins

To further verify that our identified structures are autophagy specific, we applied the same co-localization analysis for Atg14, WIPI2, Atg16L1, GEC1, and LC3B, which are involved in the formation of autophagosomes. As in the previous experiments with Atg13, those proteins were expressed as a fusion with HaloTag in HeLa cells endogenously expressing mEos2-ULK1. First, we quantified the number of structures per cell that contained more than 30 ULK1 molecules. In fed cells, either no or only a very small number of structures was detected regardless of the HaloTag-tagged autophagy proteins (Fig. 6a). Under starvation, a significant number of structures per cell with more than 30 ULK1 molecules was detected and almost all these structures contained Atg14, WIPI2, Atg16L1, GEC1, and LC3B. This result further suggests that our identified structures with more than 30 ULK1 molecules are involved in the initiation and the formation of autophagosomes. While Atg14, Atg16L1, WIPI2, GEC1, and LC3B associate with different stages of autophagosome formation, the single molecule sensitivity of our experiments revealed that they are all at least to some extent associated with the stage in which structures with more than 30 ULK1 exist.

**Fig. 6.**
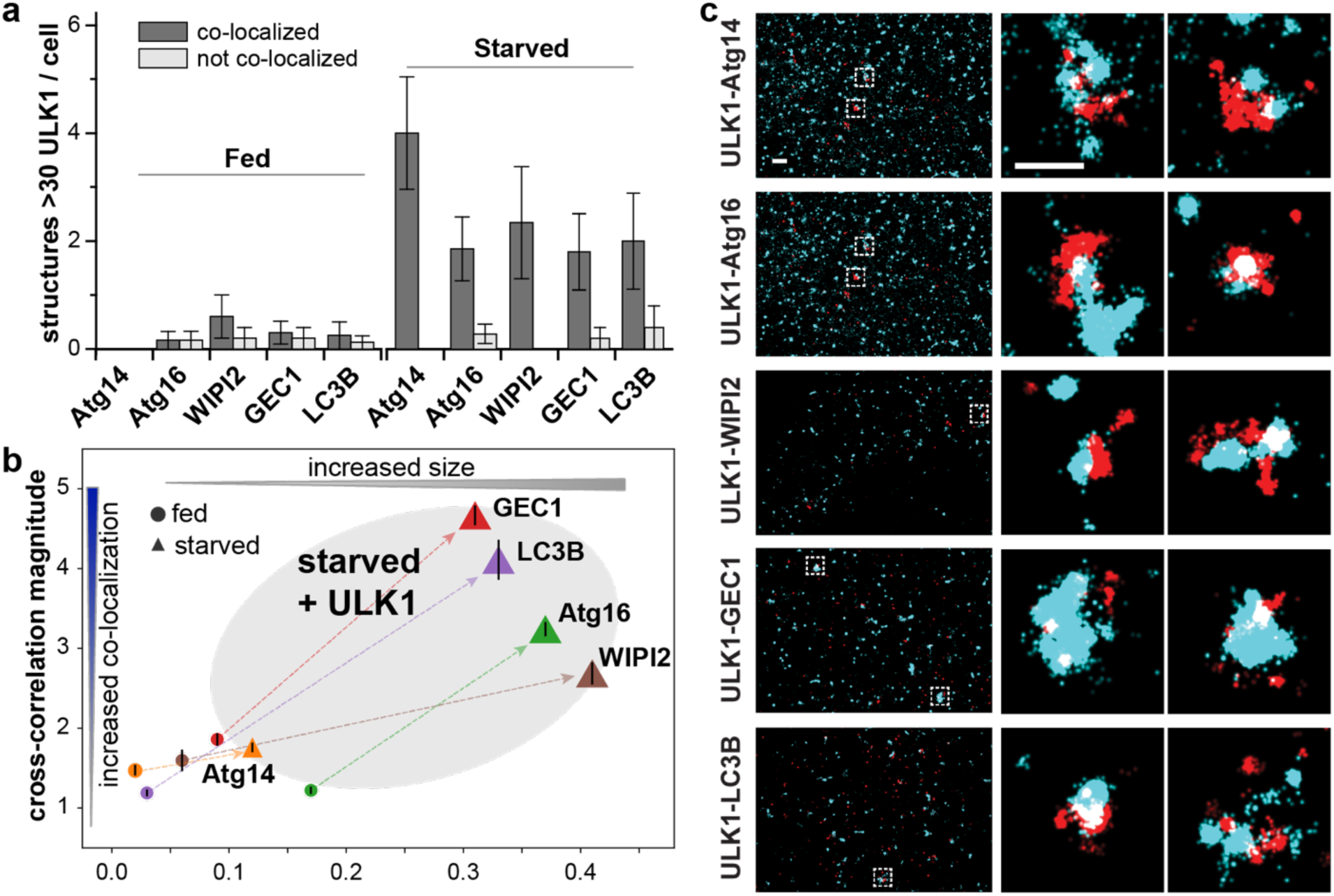
Co-localization map of ULK1 with proteins involved in autophagy on starvation-induced structures. **a** Average number of structures per cell with more than 30 ULK1 proteins that do and do not co-localize with each indicated protein. Only upon starvation, a significant number of structures with more than 30 ULK1 proteins is detected that co-localize with all proteins. **b** The cross-correlation magnitude of ULK1 with each indicated protein quantifies the degree of their overall co-localization with ULK1 in fed and starved conditions (value of 1 means no co-localization). The cross-correlation distance represents the average size of structures on which the co-localization occurs. Arrows connect the data points of each pair of proteins in fed and starved conditions. In fed cells, Atg16 and LC3B have no co-localization with ULK1 whereas Atg13, Atg14, WIPI2 and GEC1 are to some extent assembled in clusters with ULK1. All pair of proteins exhibit a significant increase of co-localization on starvation-induced structures with a significant increase in size. **c** Representative two-color PALM images under starvation and zooms that show starvation induced structures with more than 30 ULK1 molecules (red) that co-localize with the other autophagy related protein (cyan). All data recorded after 60 minutes of starvation.

To gain further insights into the nanoscopic distribution and degree of co-localization of ULK1 with those five proteins, we calculated the cross-correlation functions and quantified the peak amplitude above baseline as well as its width (Fig. 6b, Supplementary Fig. 9 and Methods). The cross-correlation magnitude above a value of one indicates the degree of co-localization and the width of the cross-correlation peak indicates the average size of structures to which the two proteins localize. In fed cells, Atg16L1 and LC3B did not co-localize with ULK1 at all, whereas Atg14, WIPI2 and GEC1, like Atg13 (Fig. 5c), were to a small degree already co-localized with ULK1. Representative two color PALM images and examples of starvation induced structures are shown in Fig. 6c. This result suggests that Atg14, WIPI2 and GEC1 are to some extent pre-assembled with ULK1 in fed cells.

Upon starvation, all proteins except Atg14 significantly increased their co-localization with ULK1. In addition, the sizes of structures to which all proteins localized significantly also increased under starvation (calculated from the width of the cross-correlation peak in Supplementary Fig. 9), consistent with the growth of autophagosomal structures. If autophagosomes form by continuous growth, as also shown by the increase of ULK1 structure sizes over time (Supplementary Fig. 7), the average size of structures might represent stages in their maturation. Consequently, we interpret that Atg13 and Atg14, which form the smallest structures, are involved in an earlier stage of autophagosome formation. WIPI2, GEC1, LC3B and Atg16 form the largest structures, which suggests their association with a later stage of autophagosome formation. Some of these results can be compared to and are consistent with previous conventional fluorescence microscopy studies^17, 18, 53^. For instance, time-lapse microscopy revealed that Atg13 localizes with LC3B to omegasomes but dissociates earlier than LC3B^18^. Consistent with this, our study showed on average smaller ULK1-Atg13 structures compared to ULK1-LC3B. Likewise, Atg14 was described to participate early in phagophore formation before Atg16L1 and LC3B become involved^17, 18^ which is confirmed by the small size of our observed ULK1-Atg14 structures compared to ULK1-Atg16L1 and ULK1-LC3B. Importantly, when comparing the cross-correlation function of structures containing more and less than 30 ULK1 molecules, it becomes apparent that co-localization of ULK1 with all other proteins mostly occurs on starvation induced structures with more than 30 ULK1 molecules (Supplementary Fig. 9). This result further highlights the importance of our identified critical number of 30 ULK1 molecules in the initiation and formation of autophagosomes and demonstrates that more than 30 ULK1 molecules are involved in all stages of autophagosome formation to which all other proteins localize.

### Vps34 and ULK1 kinase activities differentially affect ULK1 clustering during autophagy initiation

Having identified and characterized the starvation-induced structures, we wondered how the kinase activities of Vps34 and ULK1 are involved in the formation of these structures. We first treated mEos2-ULK1 HeLa cells with Vps34 inhibitor SAR405^54^, which has been shown before to block autophagy initiation and flux. When cells were starved in the presence of SAR405, the observed number of ULK1 structures per cell with more than 30 ULK1 molecules was significantly decreased by a factor of ∼4 (Fig.7a) compared to starved cells without SAR405 and was statistically similar to fed cells. The few detected clusters with more than 30 ULK1 molecules had similar sizes in the presence and absence of SAR405 (Fig.7b). These results demonstrate that Vps34 activity is required to assemble a threshold number of 30 ULK1 molecules. Since autophagy flux is also blocked as shown by the significantly reduced number of LC3B puncta per cell in the presence of SAR405 (Fig. 7c), these results highlight the importance of the critical number of ULK1 molecules to initiate autophagy.

**Fig. 7.**
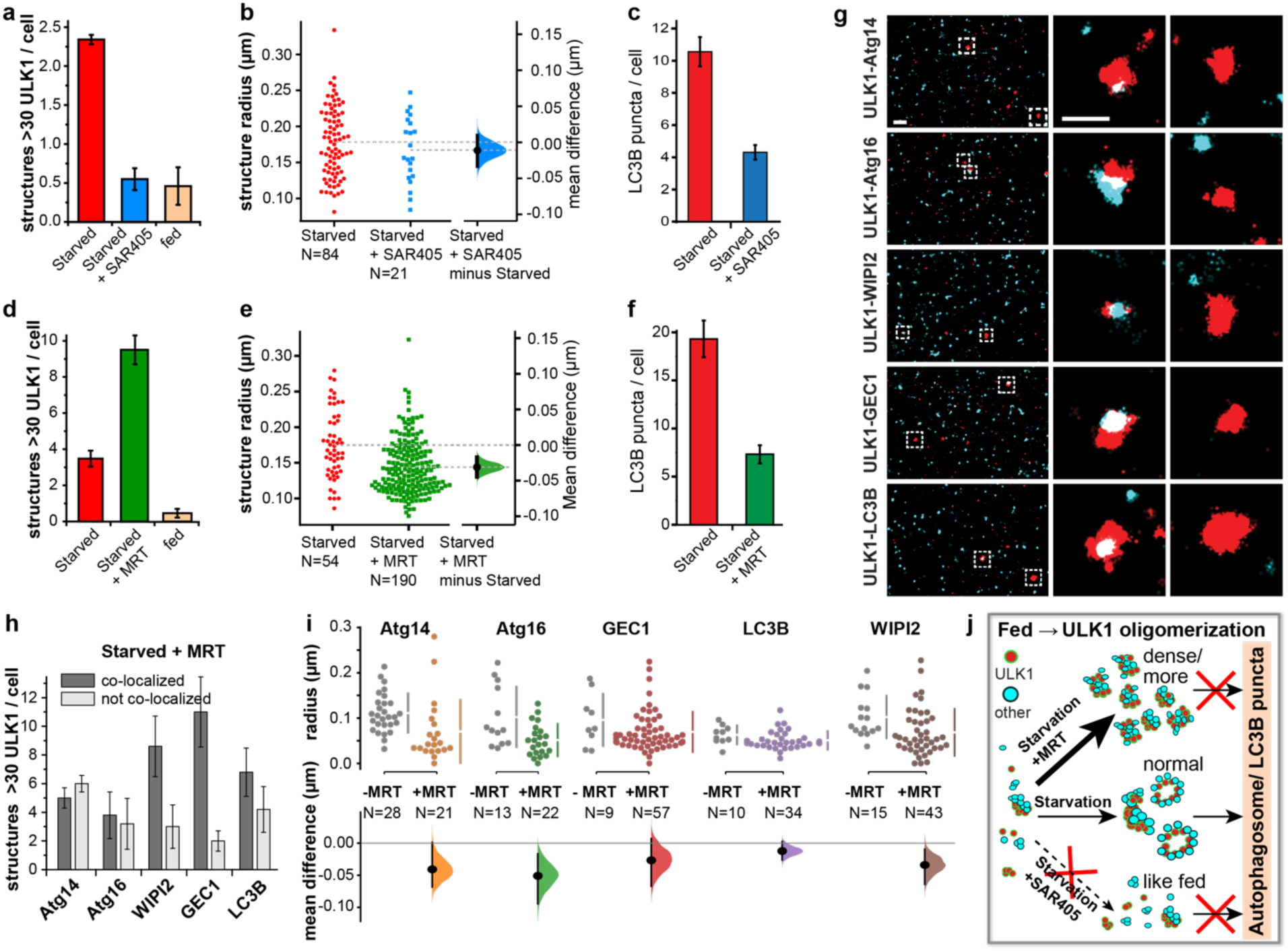
Vps34 activity is required to assemble starved-induced ULK1 structures with a threshold number of 30 ULK1 molecules, whereas ULK1 kinase activity is required for the expansion and maturation of ULK1 clusters. **a** The quantified number of structures with more than 30 ULK1 per cell is significantly decreased in starved cells in the presence of the VPS34 inhibitor SAR405 and indistinguishable from fed cells (fed: N=20, starved: N=35 starved + SAR405: N=38 cells, 150 min starvation). **b** Structures with more than 30 ULK1 molecules have no significant difference in size in the presence and absence of VPS34 inhibitor. **c** Autophagy flux as quantified by the number of detected LC3B-GFP puncta per cell is decreased in starved cells in the presence of SAR405 (N=30 cells for each condition). **d** The quantified number of structures with more than 30 ULK1 per cell is significantly increased in starved cells in the presence of the ULK1 inhibitor MRT68921 for 1h (fed: N=20, starved N=15, starved + MRT: N=20 cells, 60 min starvation). **e** The radius of structures with more than 30 ULK1 molecules is significantly smaller in starved cells in the presence of the ULK1 inhibitor (Mann-Whitney, p = 1.2×10^-5^ from 5000 bootstrap re-samplings; ANOVA, p = 1.2×10^-6^). **f** Autophagy flux as quantified by the number of detected LC3B-GFP puncta per cell is decreased in starved cells in the presence of MRT68921 (N=30 cells for each conditions). MRT68921 was added to fed cells for 30 min and then cells were starved in amino acid deficient media in the presence of the inhibitor 60 min. These results were compared to cells after 60 min of amino acid starvation. **g** Representative two color PALM images (left) and zooms of individual structures with more than 30 ULK1 molecules (right) that do and do not co-localize with other autophagy related proteins in the presence of the ULK1 inhibitor in starved cells. **h** Average number of structures per cell with more than 30 ULK1 proteins that do and do not co-localize with each indicated protein in starved cells treated with MRT68921. Significantly more structures do not co-localize with autophagy related proteins compared to starved cells in the absence of ULK1 inhibitor (N=5-8 cells for each condition, 60 min starvation). **i** The radii of structures with more than 30 ULK1 molecules measured with HaloTag localizations from each autophagy protein in starved cells is significantly smaller in the presence of MRT. **j** Schematics of ULK1 cluster formation, association with other autophagy proteins, structure formation and progression to autophagosomes for the three different conditions.

Interestingly, treating mEos2-ULK1 HeLa cells with ULK1 inhibitor MRT68921^55^ significantly increased the number of ULK1 clusters having more than 30 molecules in starved cells (3.5 ± 0.4 compared to 9.5 ± 0.8 per cell) (Fig. 7d). However, the mean cluster radius in the presence of the inhibitor (0.143 µm) was significantly smaller than that in the absence of the inhibitor (0.175 µm) (Fig. 7e), which suggests that ULK1 kinase activity is required for the expansion and maturation of ULK1 clusters. This notion is corroborated by the suppressive effect of ULK1 inhibition on LC3B puncta formation (Fig. 7f) as well as the increase of ULK1 structure sizes over time after starvation in the absence of ULK1 inhibitor (Supplementary Fig. 7).

To clarify if ULK1 inhibition not only affects the growth but also the co-localization of other autophagy related proteins, we performed PALM co-localization analysis between ULK1 and other autophagy proteins (Atg14, Atg16, WIPI2, GEC1, and LC3B) in starved cells in the presence of MRT68921. As expected, the total number of structures with more than 30 ULK1 molecules was again significantly larger in the presence ULK1 inhibitor with some variability between the different autophagy proteins due to low number statistics and cell-cell variability. However, a significantly increased number of the structures with more than 30 ULK1 molecules did not contain the other autophagy proteins compared to starved cells without the ULK1 inhibitor (Fig. 7h and Fig. 6a). The strongest effect was seen for Atg14, Atg16 and LC3B. We also analyzed the radii of Halotag localizations from the different autophagy proteins in the ULK1 structures and compared them to starved cells in the absence of the ULK1 inhibitor (Fig. 7i). In all cases, the ULK1 inhibitor significantly reduced the sizes of the structures. When comparing the cross-correlation of ULK1 and the other proteins in starved cells in the presence of the inhibitor, no co-localization was detected in structures with less than 30 ULK1 molecules as similarly as observed in the absence of the inhibitor (Supplementary Fig. 9). In structures with more than 30 ULK1 molecules, a decreased magnitude or decreased co-localization was observed for all the autophagy proteins except GEC1, which surprisingly exhibited a strongly increased co-localization. This could partially be due to the coincidentally largest number of observed structures with more than 30 ULK1 molecules per cell (Fig. 7h) or to an unknown mechanism of ULK1 activity to inhibit its association with GEC1.

These findings show that stalled autophagosomal structures with more than 30 ULK1 molecules in the presence of ULK1 inhibitor had a significantly smaller size and did not contain Atg14, Atg16, WIPI2, and LC3B to the same extent as in cells without ULK1 inhibitor. ULK1 kinase activity is therefore required for the recruitment of those autophagy regulators and for the progression and expansion of early autophagosomal structures. These results are consistent with other studies showing that ULK1 inhibition results in accumulation of stalled early autophagosomal structures^55, 56^ (Fig. 7j), and also support our finding that ULK1 is not only involved in the induction of autophagy but also in the expansion and growth of the early autophagosomal structures.

## Discussion

In this study, we have applied qPALM to quantitatively analyze the oligomeric state and the nanoscopic spatial distribution of ULK1 during autophagy initiation. By counting the number of ULK1 molecules in clusters, we found that more than 98% of ULK1 clusters (accounting for 93% of ULK1 molecules is fed and 89% in starve cells) contain 10 or less ULK1 molecules. Starvation induced about 5% of all ULK1 molecules to transition to higher order oligomeric clusters and larger structures with a significantly increased number of ULK1 molecules compared to fed cells. A small fraction of these structures contained a high number up to 161 ULK1 molecules. Only during starvation, ULK1 formed small dense clusters as well as arc and spherical structures with more than 30 ULK1 molecules that colocalized with Atg13 (Figs. 2, 4 and 5). The statistically most significant difference in the number of ULK1 molecules in clusters between fed and starved cells was 30. We therefore argue that a minimal threshold number of 30 ULK1 molecules must assemble to initiate and form autophagosomal structures.

Our quantitative PALM experiments achieved the single-molecule sensitivity with ∼25 nm spatial resolution. Although qPALM has been improved since its first development, the ability to quantify the number of molecules has remained limited. One approach for qPALM is to group localizations from the same fluorophore within a spatio-temporal threshold^32, 33, 57^. A challenge for this quantification is that PSFPs do not emit at a constant intensity but rather exhibit a variable number of fluorescent bursts separated by a variable time. This phenomenon causes single PSFPs to appear as clusters with a variable number of localizations^32, 39, 58, 59^. To quantify the oligomeric states of a protein, the PSFP must irreversibly bleach within a short time compared to the entire imaging sequence. The distribution of dark times between fluorescent bursts of single PSFP must also be known. Since the photophysical properties of PSFPs depend on the experimental conditions, we determined both spatio- and temporal thresholds for grouping localizations under our experimental condition. By performing PALM experiments at a very low photoactivation power to avoid the spatio-temporal overlap of PSFPs, we determined a dark-time cutoff of 4.5 s, which correctly groups localization of more than 99% of ULK1 molecules (Supplementary Fig. 3). In addition, we analyzed endogenously tagged ULK1 to avoid any artefact from overexpression. If a pair of PSFPs is developed suitable for molecule counting, an extension of our study could be to endogenously tag the other autophagy proteins and characterize their oligomeric states and how they relate to ULK1.

Our two-color colocalization qPALM analysis also revealed that ULK1 molecules in fed and starved cells are predominantly localized on or in proximity to the ER. While this is consistent with the broadly conceived notion that autophagosomes start to form at the ER membrane^18, 60^, our study is unique in demonstrating the dynamic alteration of ULK1 oligomeric states during the initiation process in the proximity of the ER membrane. Importantly, among the different forms of ULK1 structures, the starvation-induced structures containing > 30 ULK1 molecules were always and only found on or near the ER (Fig. 3). However, we cannot exclude the possibility that other organelles, such as mitochondria, might have contacts with those ULK1 structures.

The observed starvation-induced arc and spherical structures of ULK1 with more than 30 ULK1 molecules might present the early phagophore structures as described in a recent study by Karanasios et al.^16^. The arc-shaped structures might emerge from dense, small clusters containing both ULK1 and Atg13 and then expand to larger spherical structures. Importantly, these structures with more than 30 ULK1 molecules also contained Atg14, Atg16, WIPI2, GEC1 and LC3B, which are all known to be involved in autophagosome formation. These proteins only co-localized with the structures containing more than 30 ULK1 molecules, which further highlights the importance of this threshold number. Assuming a continuous growth of autophagosomes, the average size of structures may represent different stages in their maturation, which is also consistent with our observed increase in their size after starvation (Supplementary Fig. 7) and the increased size of structures that co-localize with the other autophagy proteins (Figs. 5, 6 and Supplementary Fig. 9).

Defining the critical number of ULK1 molecules also helped to understand the role of ULK1 and VPS34 kinase activity in the initiation of autophagy. The inhibition of VPS34 kinase activity prevented the assembly of ULK1 clusters containing more than 30 molecules. In contrast, the inhibition of ULK1 kinase activity resulted in an increased number of stalled autophagosomal structures with more than 30 ULK1 molecules (Fig. 7j). These results highlight the interwoven regulation of autophagy initiation at the level of ULK1 clustering, in which ULK1 activity acts on the expansion of autophagosomal structures, whereas VPS34 can regulate ULK1 clustering that is crucial for the initiation of autophagosome formation. ULK1 inhibition significantly increased the number of structures with more than 30 ULK1 molecule in starved cells. These ULK1 clusters were smaller in size compared to those in starved cells with ULK1 inhibition, and did not contain other autophagy related proteins to the same extent as in cells without inhibitor. This result is consistent with the recent report by Zachari et al.^56^ that ULK1 inhibitor does not inhibit the formation of ULK1 puncta in starved cells. This finding indicates that ULK1 kinase activity is not required to assemble the critical number of 30 ULK1 molecules into an autophagy initiation complex but required to drive the expansion and progression into autophagosomes.

Although our study has provided unprecedented insights into the nature and the behavior of ULK1 molecules at the initiation of autophagosome formation, it remains to understand the molecular basis for how such large ULK1 clusters form and dynamically change their multimeric state and morphology. The process may involve post-translational modifications, such as phosphorylation and acetylation, and protein-protein interactions. Combining the single-molecule qPALM analysis with biochemical techniques may be essential for achieving a more comprehensive understanding of the complex mechanisms involved in autophagy initiation.

## Methods

### Generation of genome-edited cells

To introduce the mEos2 tag into the ULK1 locus of the genome of HeLa cells, we used the CRISPR-Cas9-assisted genome editing technique^61^. The detail of the general procedure has been described in our previous report^62^. The gRNA sequence (GCGGCCGGGCTCCATGGCGC) was cloned into pSpCas9(BB)-2A-GFP (Addgene, PX458; deposited by Dr. Feng Zhang). The replacement DNA was designed to contain 700 bps of the homologous region for both sides of the inserted mEos2 sequence. Four tandem repeats of GGT nucleotide sequences were added to the end of the mEos2 sequence. The gRNA plasmid and the replacement plasmid were introduced into HeLa cells using the Neon transfection system (ThermoFisher Scientific). GFP-positive cells were sorted and plated in 96-well plates as a single cell. Individual clones were screened for expression of mEos2-ULK1 by western blotting and confirmed by genomic DNA sequencing.

### DNA constructs, stable cell generation, and transient transfection

The full length cDNAs for Atg13, Atg14, GEC1 and LC3B were described in our previous reports^6, 19, 63^. The cDNA clone for Sec61β (#49155) was obtained from Addgene, whereas the cDNA clones for Atg16L1 (BC013411) and WIPI2 (BC021200) were obtained from Open Biosystems (Huntsville, AL). The cDNA clones were amplified by polymerase chain reaction and subcloned into pLV-EF1a-IRES plasmid-blast vector (Addgene #85133), which we further engineered to introduce the HaloTag sequence for N-terminal tagging. To stably introduce the plasmids into the genome edited mEos2-ULK1 HeLa cells, we prepared lentivirus. The procedures for lentivirus preparation and target cell infection have been described in our previous report^6^. Briefly, lentivirus was prepared by transfecting HEK293FT cells (ThermoFisher Scientific, R70007) with each of the plasmids prepared as above together with pHR’8.2DR and pCMV-VSV-G DNAs using Lipofectamine 3000 (Life Technologies Scientific, L3000015) at 1:1:1 ratio. Culture medium containing virus was collected 48-60 h after transfection and filtered through 0.45 µm syringe filter (Genesee Scientific, 25-245). The target cells (mEos2-ULK1 HeLa) were infected with the collected viruses in the presence of polybrene (Sigma-Aldrich, 107689). Stably transduced cells were selected in the presence of blasticidin (8-10 µg/ml). For HaloTag-tagged Atg13 and GFP-LC3B, we transfected mEos2-ULK1 HeLa cells with prk5-HaloTag-Atg13 and pLV-mGFP-LC3B DNA using GeneJET (ThermoFisher Scientific). Twenty-four hours post-transfection, cells were used for imaging.

### Co-immunoprecipitation and western blotting

The mEos2-tagged endogenous ULK1 was enriched by immunoprecipitation using an anti-ULK1 antibody (Santa Cruz Biotech., sc-10900) following the procedure we described in our previous report ^6, 19, 62^. We used a lysis buffer containing 40 mM HEPES, pH 7.4, 120 mM NaCl, 1 mM EDTA, 50 mM NaF, 1.5 mM Na_3_VO_4_, 10 mM beta-glycerophosphate, 1% Triton X-100 (Sigma-Aldrich, X100) supplemented with protease inhibitors (Roche, 05056489001). Immunoprecipitated proteins were run on Tris-glycine gels (ThermoFisher Scientific, XP04125), transferred onto immunoblot polyvinylidene difluoride (PVDF) membranes (Bio-Rad, 1620177), and detected using ECL reagents (GenDEPOT, 20-300B). The following antibodies were used for western blotting: ULK1 (sc-33182) from Santa Cruz Biotech.; p-Atg14 Ser29 (13155) and Atg14 (96752) from Cell Signaling Technology; Atg13 antibody was described in our previous report^6^.

### Autophagy induction and flux analysis

For induction of autophagy, cells were incubated in Earle’s Balanced Salt Solution (EBSS) (Sigma-Aldrich, 2888) supplemented with 10% dialyzed fetal bovine serum (ThermoFisher Scientific, 26400044) for an indicated duration. For autophagy flux analysis by western blotting, cells were incubated in full medium or EBSS medium supplemented with serum in the presence or absence of BAFA1 (180 nM). LC3B and p62 in cell lysate were analyzed by WB using antibodies from Cell Signaling Inc (LC3B: 2775) and Santa Cruz Biotechnology (p62: sc-28359).

### Preparation of cells for imaging

HeLa cells were maintained in fluorobrite DMEM (Invitrogen) supplemented with 10% fetal bovine serum, 4 mM glutamine (Gibco), 1 mM sodium pyruvate (Gibco) and 1% penicillin-streptomycin antibiotics (Invitrogen) in a T25 flask. Cells were subcultured on eight-well chambered cover glasses (Cellvis, C8-1.58-N) 24 h before transfection or imaging. All the imaging experiments have been performed in fixed cells. For fixation, cells were treated with 3% (v/v) formaldehyde for 30 min. After fixation, we used Dulbecco’s phosphate-buffered saline (PBS) with calcium and magnesium (Gibco) for measurements. For two-color PALM imaging of mEos2-ULK1 with Atg13-HaloTag, the ER marker Sec61-HaloTag or the other HaloTag protein fusions, we used the organic dye JF 646 conjugated to the HaloTag ligands (Promega). For conventional imaging of Atg13 (Supplementary Fig. 2) Oregon green (OG) was used. Both dyes exhibit no overlap of their absorption and emission spectra with photoconverted mEos2. The dyes were added to live cells on the coverglass of the 8-well chambers at a final concentration of 500 nM for OG and 1 to 10 nM for JF 646. Cells were incubated for 15-20 min, then fixed with 3% formaldehyde and washed five times with PBS containing 1% Triton-X (TX-100) to remove excess dyes.

### PALM and conventional fluorescence data acquisition

All microscopy experiments were performed with a Nikon Ti-E inverted microscope with a Perfect Focus System and recorded on an electron-multiplying CCD camera (Ixon89Ultra DU897U; Andor) at a frame rate of 20 Hz as described in detail in the supplemental information. In short, for PALM imaging of mEos2-ULK1 one photoactivation frame (405 nm laser, 1–251 µW corresponding to a power density of roughly 0.06–15 W/cm^2^) with simultaneous transmitted light imaging was followed by nine frames of excitation (561 nm, 17 mW corresponding to a power density of roughly 1 kW/cm^2^). For two-color PALM imaging of mEos2-ULK1 and HaloTag JF646, the excitation frames consisted of five 561 nm excitation frames for mEos2 followed by five 640 nm excitation frames (640 nm, 17.5 mW, roughly 1 kW/cm^2^). Photoactivation and excitation cycles were repeated until all mEos2/JF646 molecules were imaged and bleached typically until a movie recorded at 20 Hz reached 8,000-15,000 frames. The point spread functions of single molecules were then fitted with an elliptical Gaussian function using Insight3 (Zhuang lab, http://zhuang.harvard.edu/software.html) to determine their centroid positions, intensities, widths, and ellipticities. For all localizations, the x and y coordinates, photon number, background photons, width, the frame of appearance, and other fit parameters were saved in a molecule list for further analysis. To correct for drift, all molecule lists were drift corrected using the repetitive transmitted light images ^64^. The conventional fluorescence images in Supplementary Fig. 2 were generated by averaging the 50 frames recorded at 20 Hz with a 488 nm excitation power density of 0.085 W/cm^2^, whereas in Supplementary Fig. 8 a single frame is shown. Quantification of GFP puncta per cell was achieved with Insight3 in the same way as single molecule fluorescence was quantified. Parameters for gaussian fits in Insight3 were optimized based on the sizes and intensities of puncta and consistently applied to all cells.

### Quantitative analysis of molecule numbers

Once a single mEos2 molecule is stochastically photoactivated, it emits a variable number of fluorescent bursts that are separated by a variable dark time until irreversible bleaching occurs. These fluorescent bursts can be combined since the 405 nm photoactivation rate was kept low enough to avoid spatio-temporal overlap with a nearby mEos2 molecule. The combination of these fluorescent bursts is based on two thresholds: 1) a distance threshold that specifies the maximum separation of fluorescent bursts from a mEos2 molecule (corresponding to the localization precision) and 2) a temporal dark-time threshold that specifies the maximum time between fluorescent bursts from a mEos2 molecule. The distance threshold was measured using the pair-correlation function (Supplementary Fig. 3 d), which revealed that fluorescent bursts from single mEos2 molecules are spatially separated by less than 80 nm. The dark-time threshold was determined from the cumulative dark-time histogram of fluorescent bursts and set to 4.5 seconds, which covers 99% of the dark times (Supplementary Fig. 3 e and f). Based on these two parameters, a custom written Matlab procedure combines two localizations from fluorescent bursts to a photon-weighted average position if they are separated by less than 80 nm and less than 4.5s. If a third localization again appears within the cutoff time with respect to the preceding one and within 80 nm compared to the averaged position of the previous two localizations, a new photon-weighted average position is calculated from the three localizations. This iterative process is repeated until all localizations from the same mEos2 molecules are combined to a photon weighted average position. The average position has a higher localization precision due to the increased number of photons. From the resulting blink-corrected molecule list, the detected number of mEos2 molecules and their spatial distribution can be further analyzed. It is important to note that about 40% of mEos2 molecules are not detectable as independently identified by previous studies in different model systems^32, 33^. Thus, the actual number of molecules equals the detected number divided by 0.6. Throughout this manuscript we only report the number of detected molecules for clarity.

To count the number of ULK1 molecules in initiation complexes and forming autophagosomal structures in an unbiased way, we determined their size distribution in the amino acid starved conditions using the pair-correlation function of the blink-corrected mEos2 molecule list (Fig. 2c). The pair correlation function g(r) was above 1 outside the experimental error (standard error of mean) up to a distance of 800 nm, indicating accumulation of ULK1 molecules beyond a random distribution. Therefore, ULK1 molecules within a radius of 400 nm were assigned to one structure or cluster. Structures and clusters with a certain number of ULK1 molecules were then accumulated in a histogram with bin width 10.

### Alignment of two-color PALM data

To superimpose two color PALM data for images or cross correlation analysis, we recorded conventional fluorescence images of fluorescent beads (TetraSpeck microspheres, Invitrogen T7279) that were excited at 561 nm and 640 nm and appeared in both channels. The transformation between the two channels across the field of view was then obtained by localizing individual beads in both channels and by determining the 3rd order polynomial transformation between their coordinates. The accuracy of the transformation was then verified by recording different fields of view of fluorescent beads and superimposing both channels using the determined transformation. As seen in Supplementary Fig. 10, the deviation of the bead locations from both channels was less than 19 nm, which is smaller than the localization precision of PALM.

### Spatial Cross-Correlation analysis and cluster identification

After identification of blink corrected ULK1 molecules, molecules were assigned to clusters. The radial distribution (or pair correlation) among all blink corrected molecules was calculated and molecules that were within 400 nm of each other were identified as ULK1 clusters. The obtained partial radial distribution function was normalized by the number of molecules in both datasets and the area of the field of view^39^. Next, a spatial cross-correlation between the blink corrected ULK1 molecules and localizations from the second protein of interest was performed to determine the distance distribution between the two proteins. Due to the large number of localizations in both the blink corrected ULK1 and the Atg13 localization datasets, we set a 2 µm cutoff in the nearest neighbor search, which only requires 10-30 GB of working memory. This method allows for the identification of molecule pairs within a specified distance distribution across the entire desired field of view for many movies using available computational resources (analysis code was written in MATLAB 2018b and run on an Alienware Aurora G6 computer with 3.80 GHz CPU, 40 GB of working memory, NVIDIA GeForce GTX 1070 8 GB graphics card, and 900 GB of disk space). Next, ULK1 clusters that contained at least one ULK1 molecule within 100 nm of a localization from the second protein of interest (labelled “with”) were segregated from ULK1 clusters with no molecules within 100 nm of an Atg13 localizations (labeled “without”). Cluster properties such as the radius and the number of ULK1 molecules were determined separately for a comparison of fed and starved cells. This cross-correlation analysis was also applied to ULK1 molecules and ER localizations. A detailed description of this analysis as well as the code can be found in reference^65^.

### Estimating radii of structures

To estimate the radius of the ULK1 clusters, the x and y positions of the molecules inside the cluster were averaged to find the center and the distance between each molecule and the center was calculated. The mean of the distances from the center was then used to approximate the radius of the ULK1 cluster.

### Cross-correlation distance and magnitude

The height and width of the first lobe of the cross-correlation contains inextricable information about the degree of co-localization and characteristic sizes and separation of co-localized clusters. At long distances, the cross-correlation flattens to a nearly horizontal baseline since the two molecule species are randomly distributed with respect to each other at long distances. The cross-correlation magnitude and distance in Fig. 6b was calculated by normalizing the cross-correlations, which are displayed in Supplementary Fig. 9. First, we determined a distance of 1μm was sufficiently far to achieve a flat baseline and then estimated the baseline by calculating the inverse-variance weighted mean of the cross-correlation from 1μm to 2μm. We defined the width of the cross-correlation as the farthest distance at which the cross-correlation reaches a value of 0.5 above the baseline. To determine the cross-correlation magnitude, we normalized each cross-correlation by its respective baseline so that all baselines are equal to 1 and calculated the maximum. All error bars are the standard error of the mean of the cross-correlations of individual cells, with errors propagated from the renormalization calculation.

## Acknowledgment

Research reported in this publication was supported by the National Institute of General Medical Sciences of the National Institutes of Health under Award number R21GM127965 (E.M.P) and R35GM130353 (D.H.K). We thank Yu Xu for her valuable help and input in the blink correction code. We also thank Luke Lavi’s lab and the HHMI Janelia Research campus for providing the HaloTag dye ligands.

## Notes

### Competing Interest Statement

The authors have declared no competing interest.

### Summary of Updates

The manuscript has been significantly revised to now also contain an extensive super-resolution co-localization map between ULK1 and other proteins involved in autophagy as well as a PALM analysis in the presence of ULK1 and VPS34 inhibitors.

