## Supplementary Material for "ULK1 forms distinct oligomeric states and nanoscopic structures during autophagy initiation"

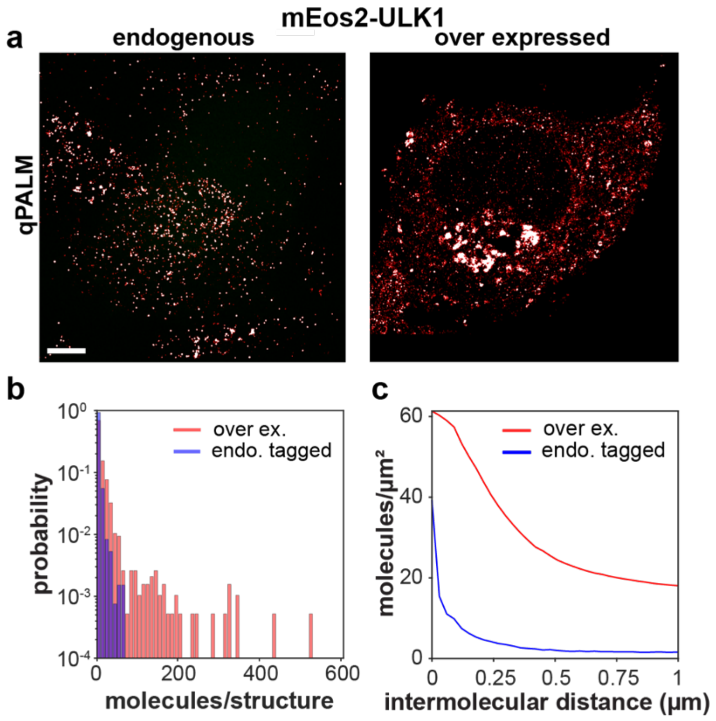


**Supplementary Fig. 1**. **Endogenous tagging of ULK1 is required to avoid artifacts in quantifying the nanoscopic distribution and oligomeric state of ULK1**. **a** qPALM images of endogenously tagged mEos2-ULK1 and overexpressed mEos2-ULK1 show that the overexpressed ULK1 forms highly dense structures, and causes clustering artifacts. Scale bar: 5 μm. **b** Analysis of qPALM data shows that overexpressed mEos2-ULK1 results in significantly more molecules and larger oligomers (red) than endogenously tagged mEos2-ULK1 (blue). ULK1-mEos2 molecules appearing within a distance of 400 nm were assigned to a structure, and a normalized histogram of the number of molecules per structure with bin width 10 was created. **c** Radial distribution comparison between overexpressed and endogenously tagged ULK1 highlights differences in the degree of ULK1 protein clustering. Overexpressed cells contain larger cluster and higher ULK1 molecule densities when compared with endogenously tagged ULK1 clusters. Condition: Amino acid starvation for 150 min.

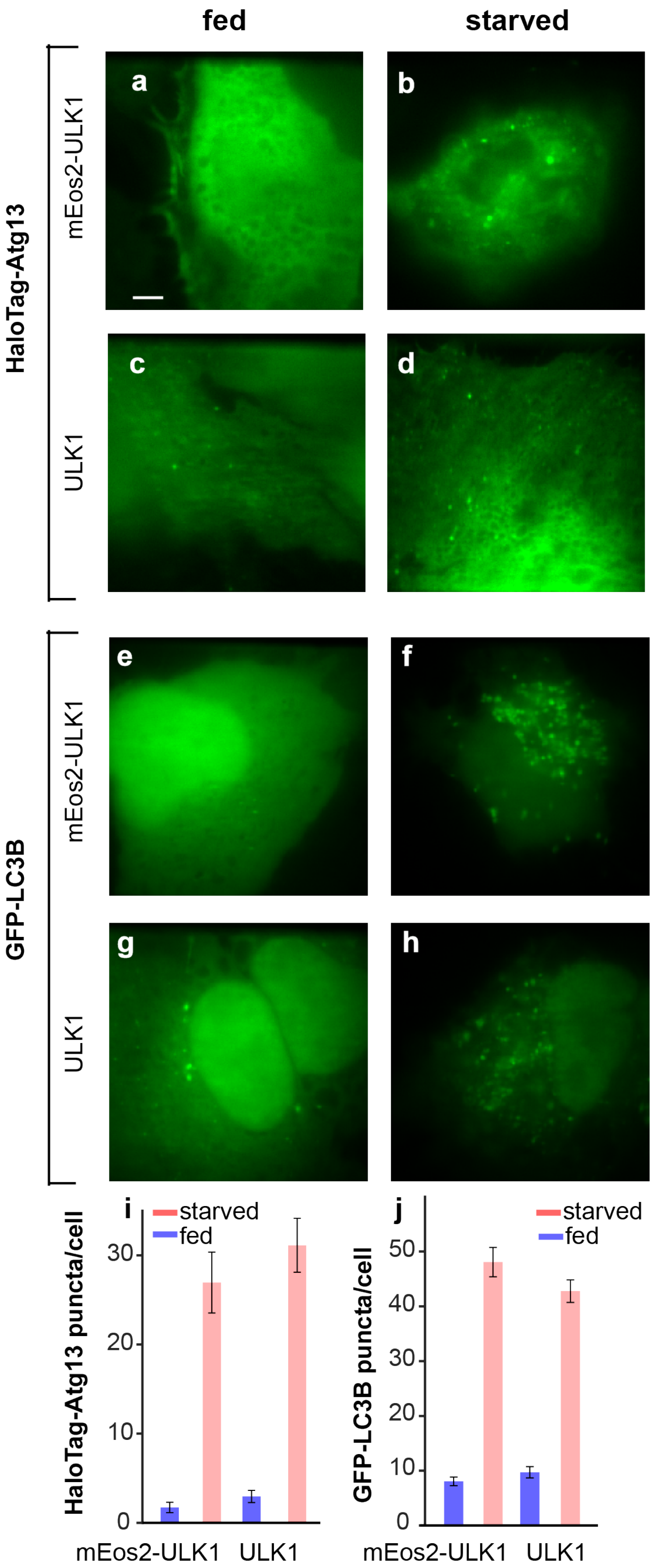
fed
starved
HaloTag-Atg13
mEos2-ULK1
ULK1
GFP-LC3B
mEos2-ULK1
ULK1
a
b
c
d
e
f
g
h
i
starved
fed
HaloTag-Atg13 puncta/cell
0
10
20
30
mEos2-ULK1
ULK1
j
starved
fed
GFP-LC3B puncta/cell
0
10
20
30
40
50
mEos2-ULK1
ULK1

**Supplementary Fig. 2. Endogenous tagging of ULK1 with mEos2 does not alter the cellular ability to form autophagosomes in response to amino acid starvation. a-d** Conventional fluorescence images of HaloTag-Atg13 bound to Oregon Green, a ligand for HaloTag. HaloTag-Atg13 was transiently expressed in unmodified HeLa cells (labeled as ULK1) or HeLa cells expressing endogenously tagged ULK1-mEos2. Cells were cultured in either full medium or amino acid-deprived medium for 150 min in the presence of BAFA1. Scale bar: 5 µm. **e-h** Conventional fluorescence images of GFP-tagged LC3B. GFP-LC3B was transiently expressed in unmodified HeLa cells or HeLa cells expressing endogenously tagged ULK1-mEos2. Cells were cultured as described in a-d. **i-j** Quantitative analysis of Atg13 and LC3B puncta from the results described in a through h. The values are means +/- standard error of the mean (number of cells analyzed from left to right: n=22, n=18, n=20, n=16, n=48, n=36, n=45, n=37). There was no statistically significant difference (ANOVA) between the modified and the unmodified cells for both LC3B and Atg13 puncta.

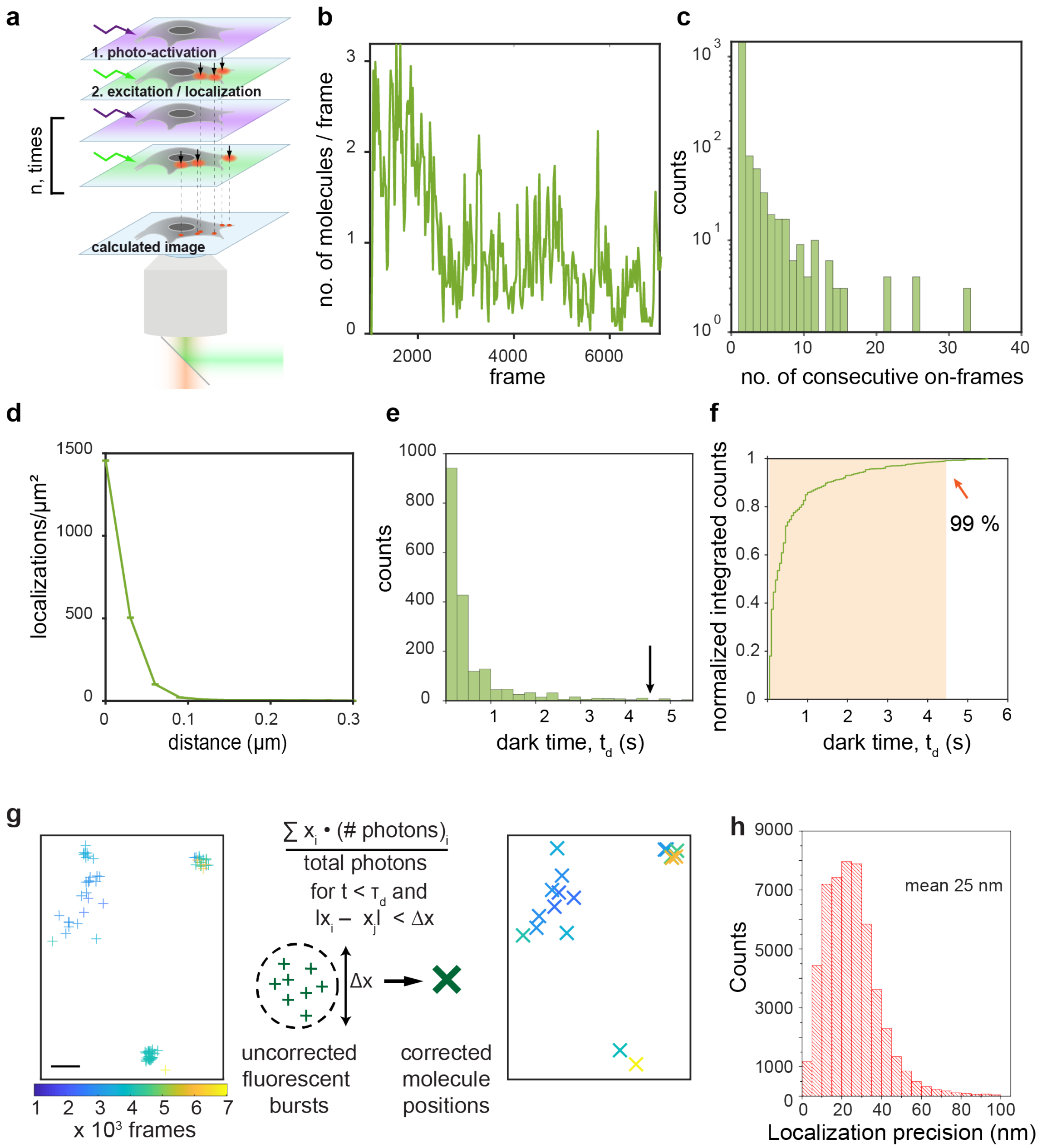


**Supplementary Fig. 3. qPALM analysis and parameters for correcting clustering artefacts from mEos2 blinking.** **a** Schematics of qPALM imaging experiments. PSFPs are stochastically activated by 405-nm light and consecutively excited by 561 nm light. Activation and excitation cycles are repeated until all molecules are imaged and bleached. Fluorescent bursts from single PSFP are localized with high accuracy by fitting their intensity profile with a gaussian function. All localizations are then superimposed and rendered in the final PALM image. **b** Number of detected mEos2-ULK1 molecules per frame in the entire field of view demonstrates that the 405 nm photoactivation power was increased slowly enough over time to avoid spatio-temporal overlap of single molecules for correct blink-correction. **c** Blinking statistics of mEos2-ULK1 molecules from a single HeLa cell showing the distribution of the number of consecutive frames in which signals from individual molecules were detected. **d** Pair correlation plot of the raw single-molecule localizations of mEos2-ULK1 from the same HeLa cell as in c. The peak width at short distances reflects the maximum distance of ~100 nm at which localizations of single mEos2-ULK1 molecules scatter as well as the clustering of mEos2-ULK1 molecules. This distance is used as the spatial cutoff for blink-correction. **e** The dark-time histogram of the ULK1-mEos2 shows the distribution of times between fluorescent bursts from individual mEos2 molecules. **f** Normalized cumulative off-time histogram and dark-time cutoff for blink correction at 4.5 s (~90 frames at 20 Hz frame rate) where 99% of fluorescent bursts are covered. **g** Left: individual single-molecule localizations are shown as crosses with a color code according to their frame of appearance. Some mEos2 molecules only appear in one frame and irreversibly bleach, whereas others blink multiple times and cause a cluster of localizations. Scale bar: 200 nm. Right: After averaging localizations appearing within the determined dark time cut-off and within the localization uncertainty, blinking of mEos2 was corrected and the number of ULK1 molecules were obtained. **h** Histogram of the localization precision of raw PALM localizations calculated with the formula by Thompson et al. based on the number of photons. The mean localization precision is 25 nm.

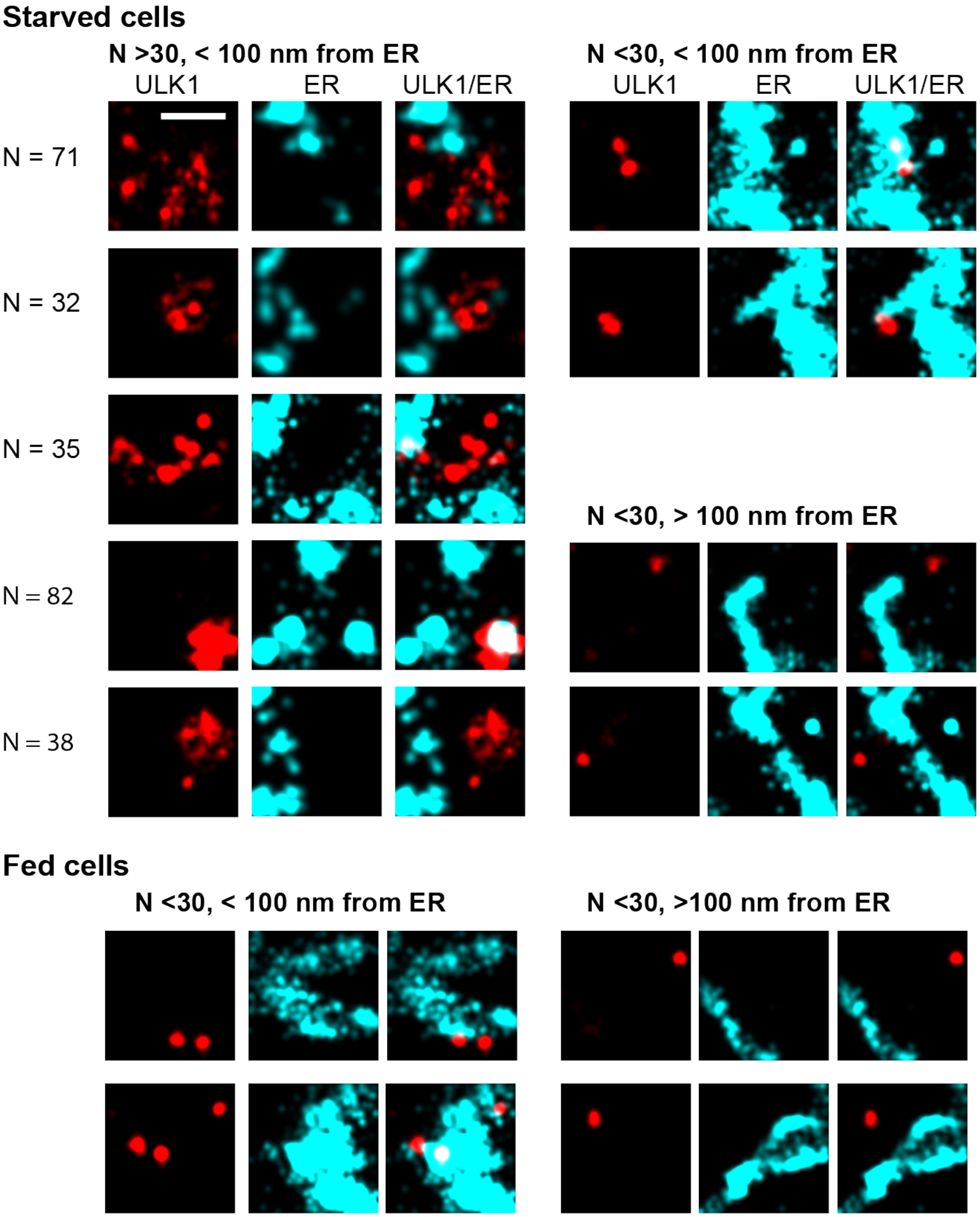


**Supplementary Fig. 4. Examples of super-resolved structures observed in fed and starved cells co-localized with ER. Upper left:** In starved cells (150 min), structures containing more than 30 ULK1 molecules are located closer than 100 nm to the ER and display a range in sizes and morphologies from large extended spherical structures to smaller spherical and arc- shaped structures. Upper right: Structures within 100 nm from the ER and less than 30 ULK1 molecules only form small but dense puncta. Structures further than 100 nm away from the ER and less than 30 ULK1 molecules only form small puncta. Bottom: In fed cells, structures that are closer than 100 nm and further away from the ER have less than 30 ULK1 molecules and form small puncta. Condition: 150 min of amino acid starvation

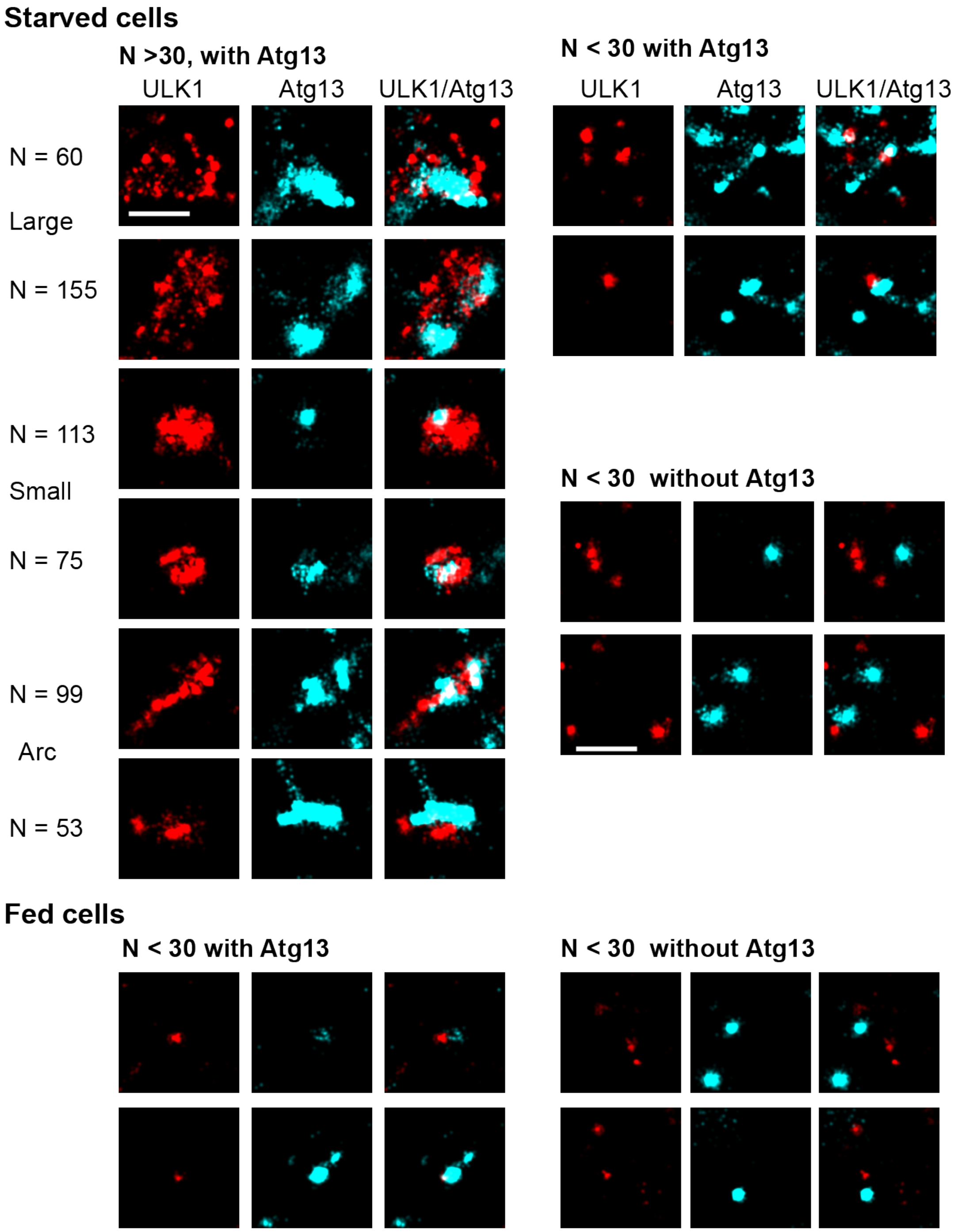


**Supplementary Fig. 5. Examples of super-resolved structures observed in fed and starved cells co-localized with Atg13. Upper left:** In starved cells, structures containing more than 30 ULK1 (red) molecules are co-localized with Atg13 (cyan) and display a range in sizes and morphologies from large extended spherical structures to smaller spherical and arc-shaped structures. Upper right: Structures with Atg13 and less than 30 ULK1 molecules only form small but dense puncta. Structures without Atg13 and less than 30 ULK1 molecules only form small puncta. **Bottom:** In fed cells, structures that do and do not contain Atg13 have less than 30 ULK1 molecules and form small puncta.

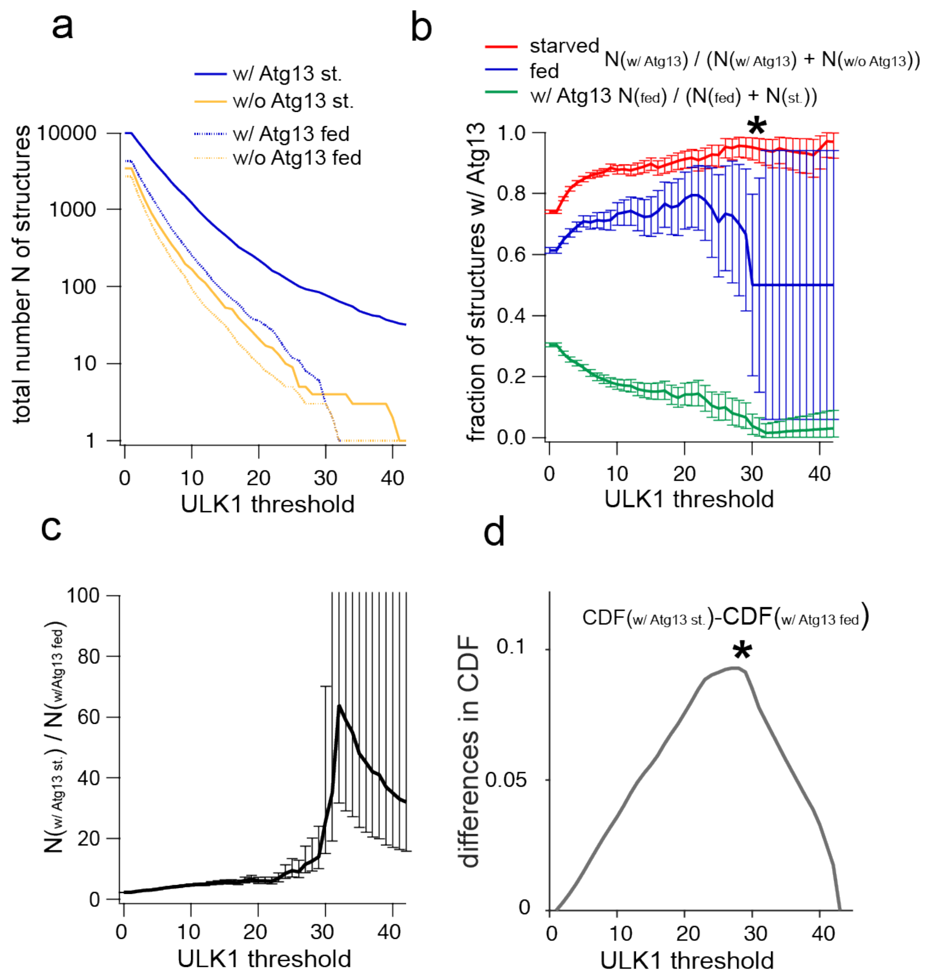


**Supplementary Fig. 6**. **Determination of a threshold for the number of ULK1 molecules in structures that do and do not contain Atg13 in fed and starved cells. a** The total number of detected structures from Fig. 5 that contain a larger number of ULK1 molecules than the respective threshold. At thresholds >43 ULK1 molecules, only Atg13-bound structures in starved cells are present. **b** Fraction of Atg13-bound structures at the respective ULK1 threshold for fed (blue) and starved cells (red). Starved cells exhibit a maximum while fed cells exhibit a minimum at a ULK1 threshold of 30 ULK1 molecules. At larger thresholds too few structures are detected in fed cells as indicated by the large error bars. The fraction of Atg13-bound structures in fed cells with respect to starved and fed cells exhibits a minimum around a threshold of 30 and errors increase at larger thresholds. Only at a threshold of 30 ULK1 molecules is the difference in the fraction between starved and fed at a maximum while the errors due to the small amount of structures detected in fed cells is small enough to yield a statistically significant difference with a p-value $<0.05$ as indicated by the asterisk (p-value from t-test). **c** Ratio of Atg13-bound structures in starved and fed cells. At a threshold of 30 the ratio is large but errors are still small enough for a meaningful comparison. At thresholds >30 ULK1 molecules too few structures are detected in fed cells, creating errors that prevent a significance testing. **d** Difference in the cumulative distribution function of fraction of Atg13 bound structures in fed and starved cells exhibits a maximum around a threshold of 30. Error bars represent the 95% confidence interval based on the absolute number of structures calculated with the inverse beta function.

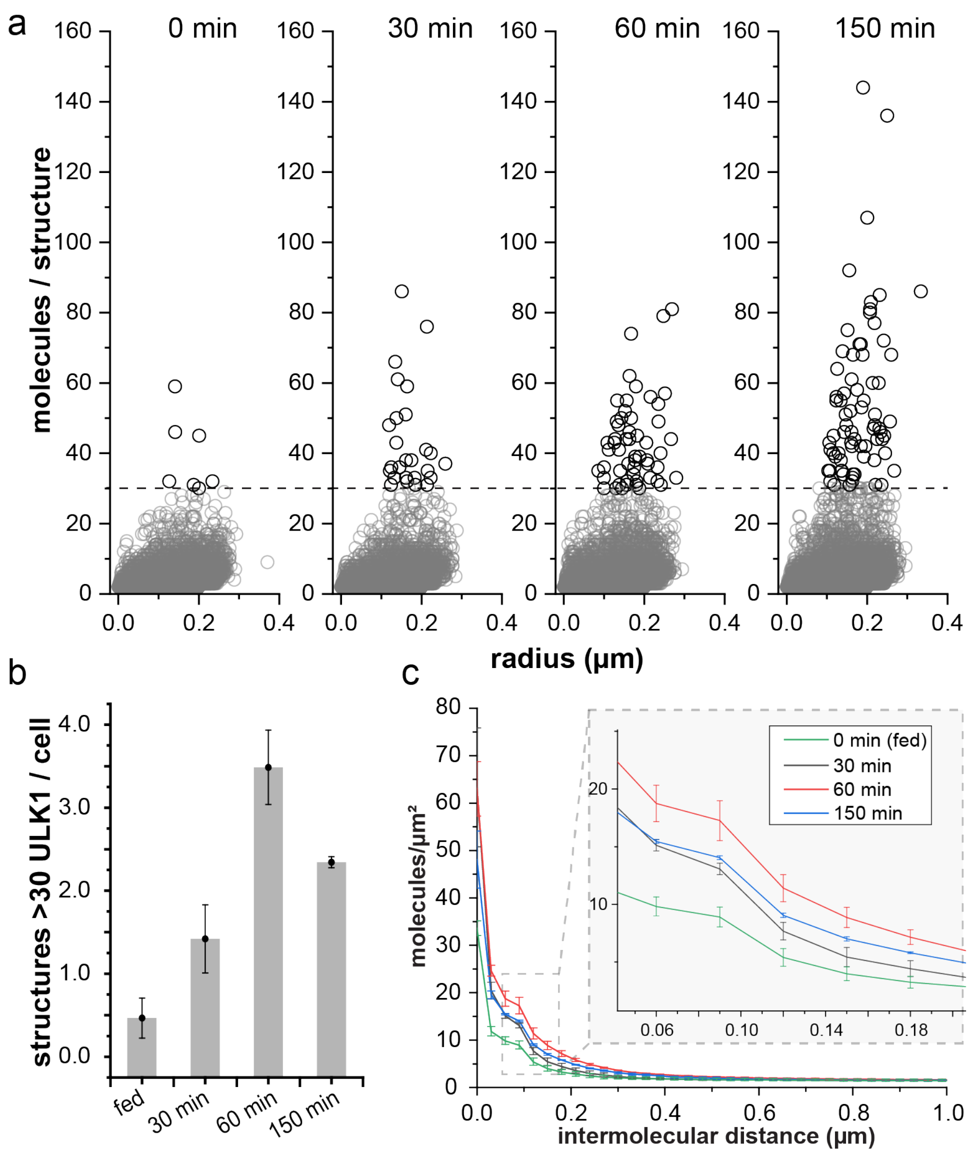


**Supplemental Fig. 7. Time-dependent clustering and ULK1 structure formation after starvation**.

**a** Scatter plot of the number of ULK1 molecules in structures vs. approximated structure radius after different starvation times. Over time, more and more structures with more than 30 ULK1 molecule are detected and the maximum number of ULK1 molecules also increases for some time points. In general, the radius of structures at early time points appears smaller whereas larger structures are detected at later time points. **b** Number of structures with more than 30 ULK1 molecules per cell. Initially a significant increase in starvation-induced structures with more than 30 ULK1 molecules is detected until 60 minutes of starvation. After 150 mins of starvation, a down-regulation of these starvation induced structures is observed presumably due to the downregulation of autophagy. **c** Pair correlation function of mEos2-ULK1 after 0 min to 150 mins of amino acid after starvation. The solid line represents the mean value of the three independent experiments and the errors the SEM. The size and density of the ULK1 clusters increased until 60 min of amino acid starvation, and then decreased at 150 min. Data from three independent experiments, total of 30 to 40 cells in each condition.

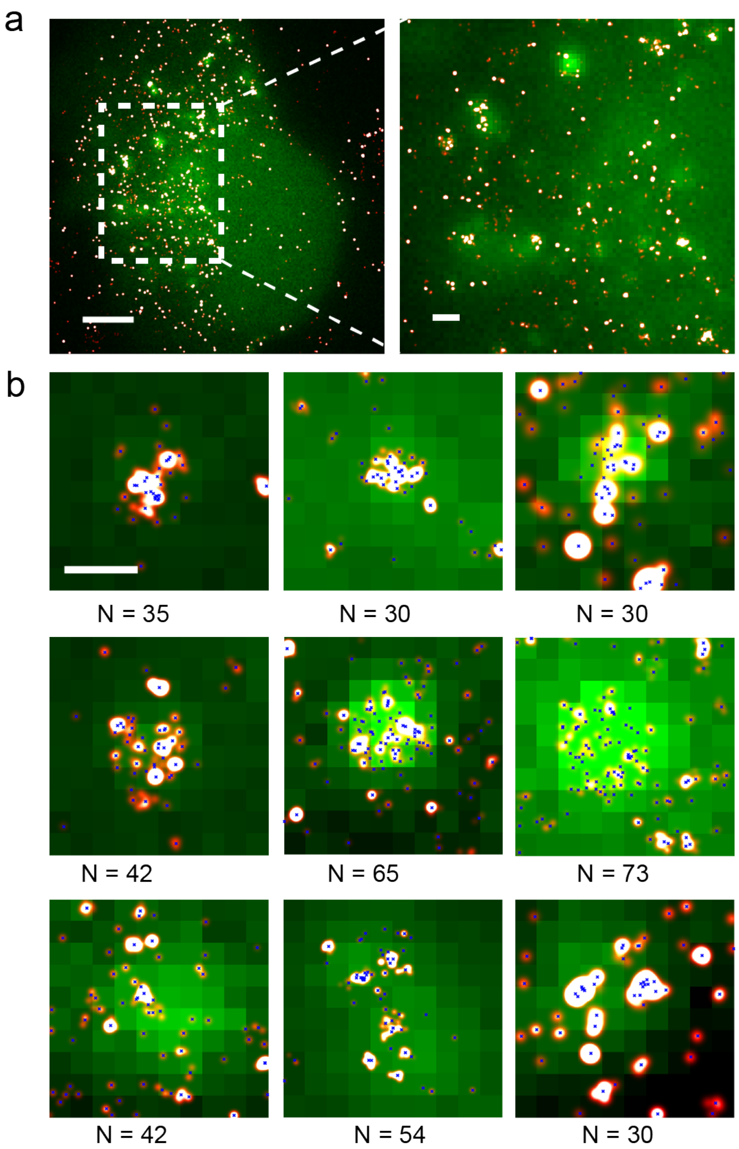


**Supplementary Fig. 8.** Co-localization of mEos2-ULK1 with the transiently transfected phagophore marker LC3B-GFP reveals that larger ULK1 structures are phagophores. **a** Two-color overlay image of mEos2-ULK1 (red) and LC3B-GFP (green) (scale bar left: 5 µm; right: 1µm). **b** The magnifications show different ULK1 structures that co-localize with LC3B, including the number of ULK1 molecules (N) they contain (scale bar 500 nm). Condition: 150 min starvation in the presence of BAFA1.

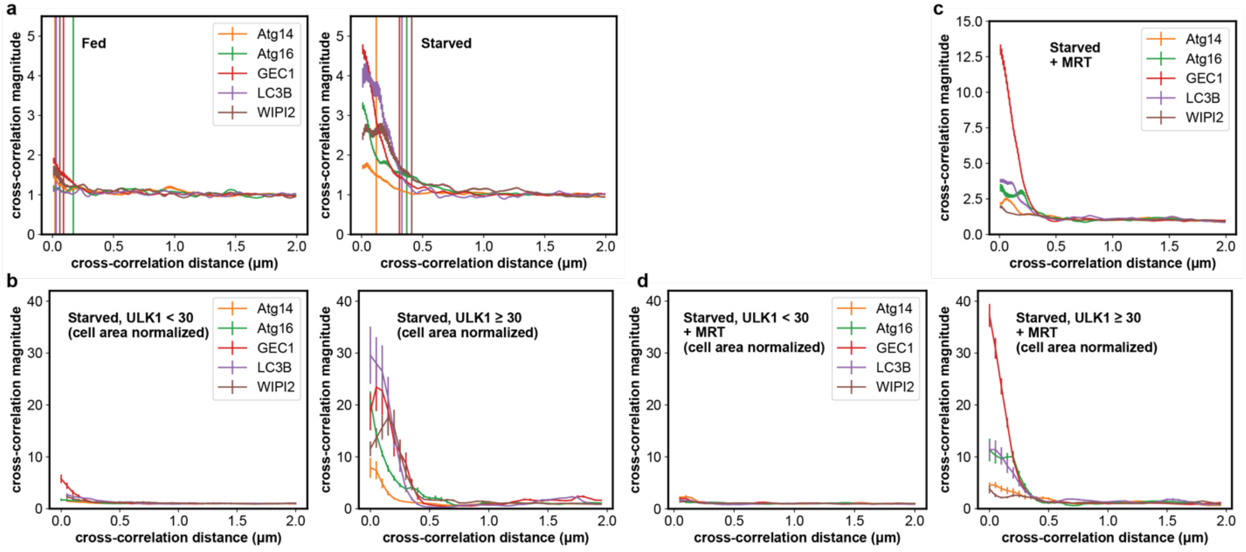


**Supplementary Fig. 9.** Cross-correlation function quantifies the degree of co-localization between ULK1 and each indicated autophagy related protein in two-color PALM data under different conditions **a** Cross-correlation between all ULK1 localizations and all localizations of the second protein pair in fed (left) and starved (right) conditions used for Fig. 6b. In fed cells, the magnitude and width of the cross-correlation is significantly smaller compared to fed cells, showing the starvation-induced co-localization to large structures. **b** Cross-correlation as in a (starved) but separately analyzed for localizations that reside in structures with less than 30 ULK1 molecules (left) and more than 30 ULK1 molecules (right). Only in structures with more than 30 ULK1 molecules, a significant peak and cross-correlation values above 1 are observed, demonstrating that co-localization mostly occurs in starvation-induced structures. **c** Cross-correlation between all ULK1 and each indicated autophagy related protein in starved cells and the presence of the ULK1 inhibitor MRT. **d** When the molecule lists are again split into structures that contain less (left) and more (right) than 30 ULK1 molecules, it becomes apparent that almost all co-localization occurs on structures with more than 30 ULK1 molecules. All cross-correlations have been normalized for better comparison as described in the Methods.

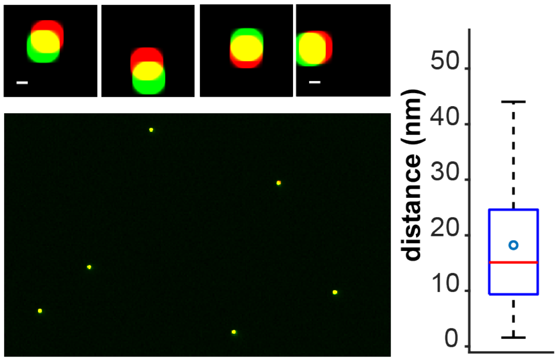


**Supplementary Fig. 10**. **Discrepancy of the transformation from the red to the green channel**. Left: Two-color overlay of the localizations of fluorescent beads (TetraSpeck microspheres, ThermoFisher T7279), that have been excited with 561 nm (green) and 640 nm (red), detected in both channels and transformed with a 3rd order polynomial to determine colocalization (upper, scale bar 20 nm) Examples of the deviations of the destination and transformed source bead positions. Right: Box plot of the distance between the same beads in the red channel and transformed green channel quantifies the precision of the two-color co-localization. Boxes extend from the 25th to 75th percentiles (interquartile range, IQR). Whiskers extend to the farthest data point within 1.5 IQR below and above the 25th and 75th percentiles, respectively. The red horizontal line represents the median accuracy of 15 nm. The cyan circle represents the mean (18 nm).
